# ROS inhibits microtubule dynamics and cell growth heterogeneity during Arabidopsis sepal morphogenesis

**DOI:** 10.1101/2025.05.09.653143

**Authors:** Isabella Burda, Fridtjof Brauns, Aaron Shipman, Emily Shapland, Lilan Hong, Adrienne Roeder

## Abstract

Developing organs grow to reproducible sizes and shapes yet the growth of their constituent cells can be highly heterogeneous and fluctuating. During wild-type *Arabidopsis thaliana* sepal development, the fluctuations in cell growth average such that the sepals grow to uniform sizes and shapes. The uniform size and shape of the sepals allow the flower bud to stay closed and protected until the floral organs are mature. In contrast, cell growth averaging is reduced in the *ftsh4-5* mutant, and the sepals develop to variable sizes and shapes. *FTSH4* encodes a mitochondrial i-AAA protease localized to the mitochondria. Reactive oxygen species (ROS) accumulate in *ftsh4-5* mutants, and lowering ROS levels rescues the sepal size and shape variability. Here, we investigate the effects of ROS on cell growth heterogeneity and cortical microtubule dynamics. We find that elevated ROS suppresses cell growth heterogeneity and averaging. We also find that elevated ROS causes cortical microtubules to become more ‘crisscrossed’, as well as more stable. The growth of the cells with crisscrossed microtubules changes less in time, which impairs cell growth averaging. However, depolymerizing microtubules is insufficient to restore normal growth fluctuations. Altogether, our results suggest that ROS affects microtubule dynamics and cell growth fluctuations which are necessary for robust morphogenesis.

## Introduction

Organ size and shape are important for function (Hong et al., 2018). For example, organs need to scale to be the correct size for the organism or have specialized shapes for a certain task. Sepals, the four leaf-like organs on the outside of the flower, have uniform size and shape both within individual flowers and between flowers of other plants (Hong et al., 2016; Zhu et al., 2020). Uniform sepal size and shape allows flower buds to remain closed while the inner floral organs are developing, protecting them from the environment. Since sepal size and shape is a reproducible outcome of development despite fluctuations in cell growth and division (Burda et al., 2024; Hong et al., 2016; Tauriello et al., 2015), this means that development is robust or not easily influenced by noise.

Even when development is robust, or reproducible, cell growth rates and directions can be heterogeneous (Elsner et al., 2012; Hong et al., 2016; Tauriello et al., 2015; Uyttewaal et al., 2012). Sometimes heterogeneity can be more effective for achieving robust morphogenesis than homogenous cell behavior. For example, during the closing of the *Xenopus laevis* neural tube, spatially random cell constriction occurs and modeling suggests this is more effective at folding the tissue than simultaneous constriction (Suzuki et al., 2017). In wild-type sepal development, epidermal cell growth rates fluctuate around an average target growth rate (Burda et al., 2024; Hong et al., 2016). Growth fluctuations are not correlated temporally or spatially in the tissue, so growth averages to the target rate over time and over larger spatial regions of the tissue (Burda et al., 2024). This means that cumulative growth is similar throughout the tissue, and thus final sepal size and shape is uniform between sepals on the same flower and between different flowers (Burda et al., 2024; Hong et al., 2016). Therefore, spatiotemporal averaging of growth heterogeneity is a mechanism for developmental robustness. Averaging of heterogeneity is also a generalizable concept; for example, noise in transcript levels has also been shown to average over time and spatially in a tissue (Little et al., 2013).

Spatiotemporal averaging is disrupted in the *ftsh4-5* mutant (Hong et al., 2016), which is a null mutant for the mitochondrial i-AAA protease FtsH4 (FILAMENTATION TEMPERATURE SENSITIVE H4) (Urantowka et al., 2005). Growth fluctuations in *ftsh4-5* are correlated in time, meaning that cells grow faster or slower than the target growth rate for too long. As a result, growth accumulates in patches of overgrowth and undergrowth, which leads to variable sepal size and shape (Burda et al., 2024).

Mutants for *ftsh4* also have abnormal mitochondria morphology with few cristae (Gibala et al., 2009) and accumulate aggregates of proteins largely composed of mitochondria small heat shock proteins (Maziak et al., 2021).This suggested that Reactive oxygen species (ROS) might be involved, as these are a normal byproduct of aerobic respiration that are toxic at high levels, and have also been co-opted as signaling molecules (Mittler, 2017). ROS are elevated in *ftsh4-5* mutants which is linked to the severity of the serrated leaf phenotype (Gibala et al., 2009). Overexpression of *CATALASE2*, which breaks down hydrogen peroxide, decreases ROS levels, rescues *ftsh4-5* sepal variability phenotype (Hong et al., 2016). Since elevated ROS causes loss of robust sepal development, we hypothesized that ROS may increase temporal correlations in growth fluctuations, which inhibits averaging to a target growth rate.

Multiple factors have been found to contribute to growth heterogeneity. In development of the Arabidopsis sepal epidermis, some of this heterogeneity derives from changes in growth rate as cells differentiate into specialized cell types (Le Gloanec et al., 2022) or from decreases in growth rate of cells neighboring the differentiating cells to buffer the fast growth (Hervieux et al., 2017; Le Gloanec et al., 2022). In the *vip3* mutant, increased noise in gene expression has been associated with increased cell growth heterogeneity and organ shape variability (Trinh et al., 2023), yet it is not known whether stochastic gene expression in wild type also drives heterogeneous cell growth. In the shoot apical meristem, mutants in a pair of actin genes have reduced cell growth heterogeneity (Wang et al., 2024).

Likewise cortical microtubules contribute to growth heterogeneity, which is reduced when microtubule dynamics are impaired in *katanin* mutant meristems (Uyttewaal et al., 2012). The mutant *mor1-1*, which also affects microtubule dynamics, slightly increases heterogeneity in subcellular growth direction near abnormal cell shapes (Elsner et al., 2025). Although some sources of heterogeneity are known, the pathways by which they create growth heterogeneity are unclear.

Plant cell growth occurs when the cell wall is deformed by turgor pressure; therefore, the mechanical properties of the cell wall could affect growth heterogeneity. Cellulose microfibrils are polysaccharide chains that are a component of the cell wall (Somerville et al., 2004), and their layout confers anisotropic mechanical properties to the cell wall (Coen and Cosgrove, 2023). The cell wall will be stiff in the direction parallel to cellulose microfibrils; as a result, turgor pressure deforms the wall perpendicular to the cellulose microfibrils (Coen and Cosgrove, 2023). Cellulose synthases move along cortical microtubules (Paradez et al., 2006), and thus the direction of new cellulose microfibrils added to the cell wall should mirror the cortical microtubule arrangement. Cortical microtubules also respond to growth heterogeneity by aligning parallel to tension generated in cells neighboring a rapidly growing cell. This causes the neighboring cells to decrease their growth rate (Hervieux et al., 2017). Therefore, a feedback loop exists because microtubule orientation is affected by cell growth and tension, and then microtubule orientation affects cell growth through the orientations of cellulose microfibrils.

Here we investigate the effect of ROS levels on spatiotemporal averaging of growth rate fluctuations. We also test whether ROS affects microtubule dynamics. We find that elevated ROS levels inhibit growth heterogeneity and increase microtubule stability. The cells with decreased microtubule dynamics also display temporally correlated growth, although depolymerizing the microtubules is insufficient to restore normal growth fluctuations. Our results show that ROS homeostasis, microtubule dynamics, and low correlations in cell growth fluctuations contribute to robust morphogenesis.

## Results

### ROS inhibits cell growth heterogeneity

The loss of developmental robustness mutant, *ftsh4-5*, was previously shown to have increased ROS and to be rescued by overexpression of *CATALSE2*, which decreases ROS by breaking down hydrogen peroxide. This suggests it is the accumulation of ROS that leads to sepal size and shape variability (Hong et al., 2016). Here, we examined *ftsh4-5*, overexpression of *CATALASE2* (*p35S::CAT2*, hereafter referred to as *CAT2oe*), *ftsh4-5 CAT2oe*, and wild type to investigate the relationship between ROS levels, microtubules, and cell growth heterogeneity during robust development. The *CAT2oe* and *ftsh4-5 CAT2oe* plants were heterozygous for individual insertions of the transgene. Independent insertions of *CAT2oe* had differing expression levels of *CATALASE2* (Fig S1A) and genotyping confirmed the transgene was segregating in the offspring. To check how *CAT2oe* affected ROS levels, inflorescences were stained for hydrogen peroxide (Fig 1A-D, Fig S1B) and superoxide (Fig 1E-H, Fig S1C) which are two species of ROS. ROS levels in wild type are low in young flower buds (Fig 1A, E), later appearing at the tip of sepals in older flowers and progressing proximally down sepals as flowers continue to develop (Hong et al., 2016) (Fig S1B-C). The spatial localization of ROS appears similar in *CAT2oe* as in wild type. *CAT2oe* has slightly lower levels of ROS, suggesting that it has only a marginal effect when ROS is already at wild type levels (Fig 1B, F, Fig S1B-C). In *ftsh4-5*, ROS is elevated in flowers of all developmental stages (Hong et al., 2016) (Fig 1C, G, Fig S1B-C). However, in *ftsh4-5 CAT2oe*, ROS is lowered in young flowers compared to *ftsh4-5* (Fig 1D, H), and its localization appears similar to the wild-type localization (Fig S1B-C). Our results confirm rescue of the *ftsh4-5* high ROS phenotype by promoting ROS breakdown as shown previously (Hong et al., 2016).

**Figure 1:**
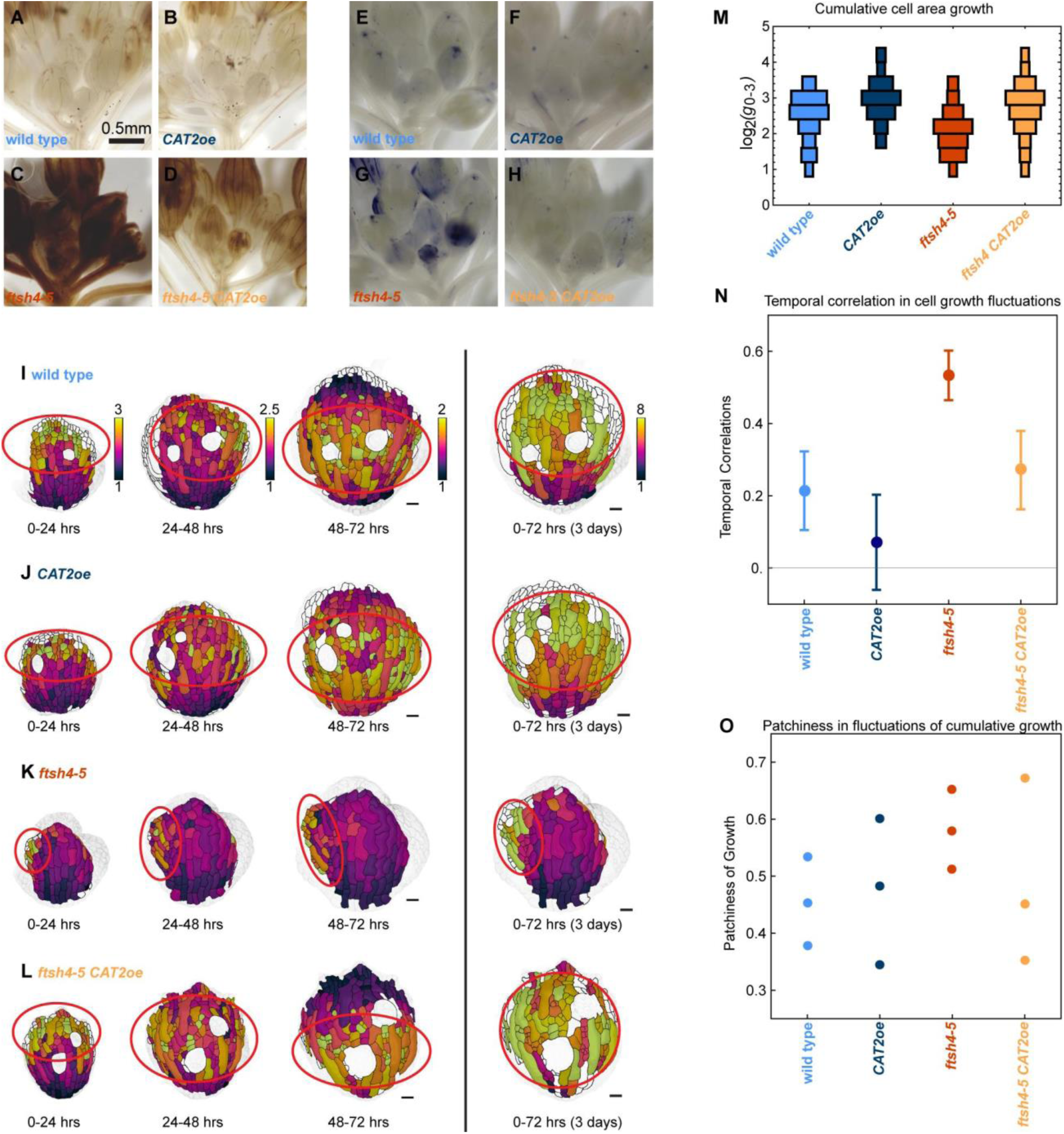
ROS inhibits growth heterogeneity. A-D: Inflorescences stained for hydrogen peroxide. Scale bars are 0.5 mm. Representative images from n=3 biological replicates. *CAT2oe* and *ftsh4-5 CAT2oe* are heterozygous for individual insertions of *CAT2oe*, and from the same plants that were live imaged. (A) wild type and (B) *CAT2oe* have similar levels of hydrogen peroxide, (C) *ftsh4-5* has elevated levels of hydrogen peroxide, and (D) *ftsh4-5 CAT2* has a partially rescued level. E-H: Inflorescences stained for superoxide. (E) wild type and (F) *CAT2oe* have similar levels of superoxide, (G) *ftsh4-5* has elevated levels of superoxide, and (H) *ftsh4-5 CAT2* has a partially rescued level. I-L: Cell area growth represented as a ratio, projected on the later time points over 24-hour intervals and cumulative over 3 days. (I) Wild type and (J) *CAT2oe* cell growth follows a basipetal (base to tip) growth pattern and local heterogeneity over 24 hr intervals, and the heterogeneity accumulates evenly over 3 days. (K) *ftsh4-5* cell growth follows a basipetal growth pattern but with growth asymmetrically localized within the organ-scale pattern, and it accumulates into patches of more or less cumulative growth over 3 days. (L) *ftsh4-5 CAT2* cell growth is rescued and is similar to wild type over 24 hr intervals and 3 days. Scale bars are 20μm. Representative images from n=3 biological replicates. (M) Histograms of area growth rate in all four genotypes (n=3 sepals) with cell growth calculated as a ratio of initial area to final area. (N) Plot of the average value of temporal correlation in growth rates for all cells imaged in a sepals (for n=3 sepals). The fluctuations in growth rate around the average growth for each time point and replicate are calculated, and then the temporal correlation in fluctuations over subsequent 24 hr intervals is calculated. To calculate growth fluctuations, the growth rate along the proximal-distal axis and fit with a third order polynomial, which is then considered the average growth. The difference between actual growth rates and the average growth rate are the fluctuations in growth. For more detail on how fluctuations were calculated, see Methods and Burda et al 2024. ** means p value <0.01, *** means p value <0.001, and **** means p value<0.0001. (O) Patchiness of growth over 3-days is calculated by the standard deviation of the fluctuations after averaging the fluctuation over the immediate neighboring cells.

To test whether ROS levels affect correlations in growth fluctuations and spatiotemporal averaging, we time-lapse live-imaged sepal development of plants expressing fluorescent markers for the plasma membrane (*pUBQ10::mCherry-RCI2A*) and microtubules (*pUBQ10::GFP-MBD*) (Figure 1I-L). To assess growth heterogeneity, we quantify the fluctuations from the average growth, and then calculate the correlation in fluctuations of each cell between time intervals (Fig 1N; as described previously in Burda et al., 2024). Wild type has low temporal correlations in growth fluctuations and so over the 3-day imaging series, fluctuations average and the cumulative cell growth rates are similar between cells (Fig 1I, Fig S1D-E). This is spatiotemporal averaging of heterogeneity (Burda et al., 2024; Hong et al., 2016). *CAT2oe* also has low temporal correlations in growth fluctuations which average in the 3-day cumulative cell growth rates (Fig 1J,N Fig S1F-G). Growth fluctuations are temporally correlated in *ftsh4-5*, with regions of cells that grow slower than average over all 24-hr time intervals, and regions of cells that grow faster throughout all 24-hr time intervals, resulting in 3-day cumulative cell growth rates that appear asymmetric and patchy (Fig 1K,N Fig S1H-I) (Burda et al., 2024). In *ftsh4-5 CAT2oe*, temporal correlations in growth fluctuations are lowered to levels similar to wild type, and the cumulative 3-day growth is averaged like wild type (Fig 1L,N Fig S1J-L), consistent with the restoration of robust sepal morphology. Additionally, cumulative growth rate is decreased in *ftsh4-5*, likely due to the contribution of slow-growing patches of cells and increased in *ftsh4-5 CAT2oe* although the difference does not reach significance (Fig 1M). Together, this indicates that *CAT2oe* restores growth heterogeneity in *ftsh4-5* as well as restoring sepal morphology.

Increased temporal correlations in growth heterogeneity leads to cumulative growth that is unevenly distributed in the organ, with patches of overgrowth and undergrowth (Burda et al., 2024). Patchiness is quantified by averaging the fluctuation in 3-day cumulative growth of each cell with its immediate neighboring cells and then calculating the variability in these averaged fluctuations (Fig 1O; as previously described in Burda et al., 2024). A lower score indicates more evenly distributed cumulative growth whereas a higher score indicates more unevenly distributed or patchy cumulative growth.

Patchiness of the 3-day growth is partially rescued in *ftsh4-5 CAT2oe*, but the differences between groups does not reach significance (Fig 1O). The difference in significance between these results and previous results (Burda et al., 2024) is likely due to the slightly slower growth of the microtubule marker line, so that the *ftsh4-5* patchy growth is not as pronounced in the same time interval. The variability in the rescue comes from natural variation in severity of the *ftsh4-5* phenotype and likely from differences in expression of *CAT2* in different transgenic lines. Together, this indicates that high ROS increases temporal correlations in growth fluctuations, inhibiting averaging of growth fluctuations.

### Increasing ROS leads to crisscrossed microtubule arrangement

Next, we investigated whether changes in cell growth heterogeneity correlated with differences in microtubule arrangement. Since the microtubule marker was imaged simultaneously with the membrane marker, we examined the microtubule arrangement of individual cells at each time point of the live imaging series from Figure 1 and S1 (Fig 2A-B, Fig S2A-J). At the beginning of the time series (corresponding to floral stage 5), microtubules in wild type appear isotropic (meaning in all directions) or slightly anisotropic in the transverse direction (parallel to the short axis of the cell) (Fig 2C). *CAT2oe* microtubules appear similar to wild type (Fig 2D). However, microtubules in *ftsh4-5* often appear longitudinal (parallel to the long axis of the cell) (Fig 2E). This is rescued in *ftsh4-5 CAT2oe* which have microtubules arranged similar to wild type (Fig 2F).

**Figure 2:**
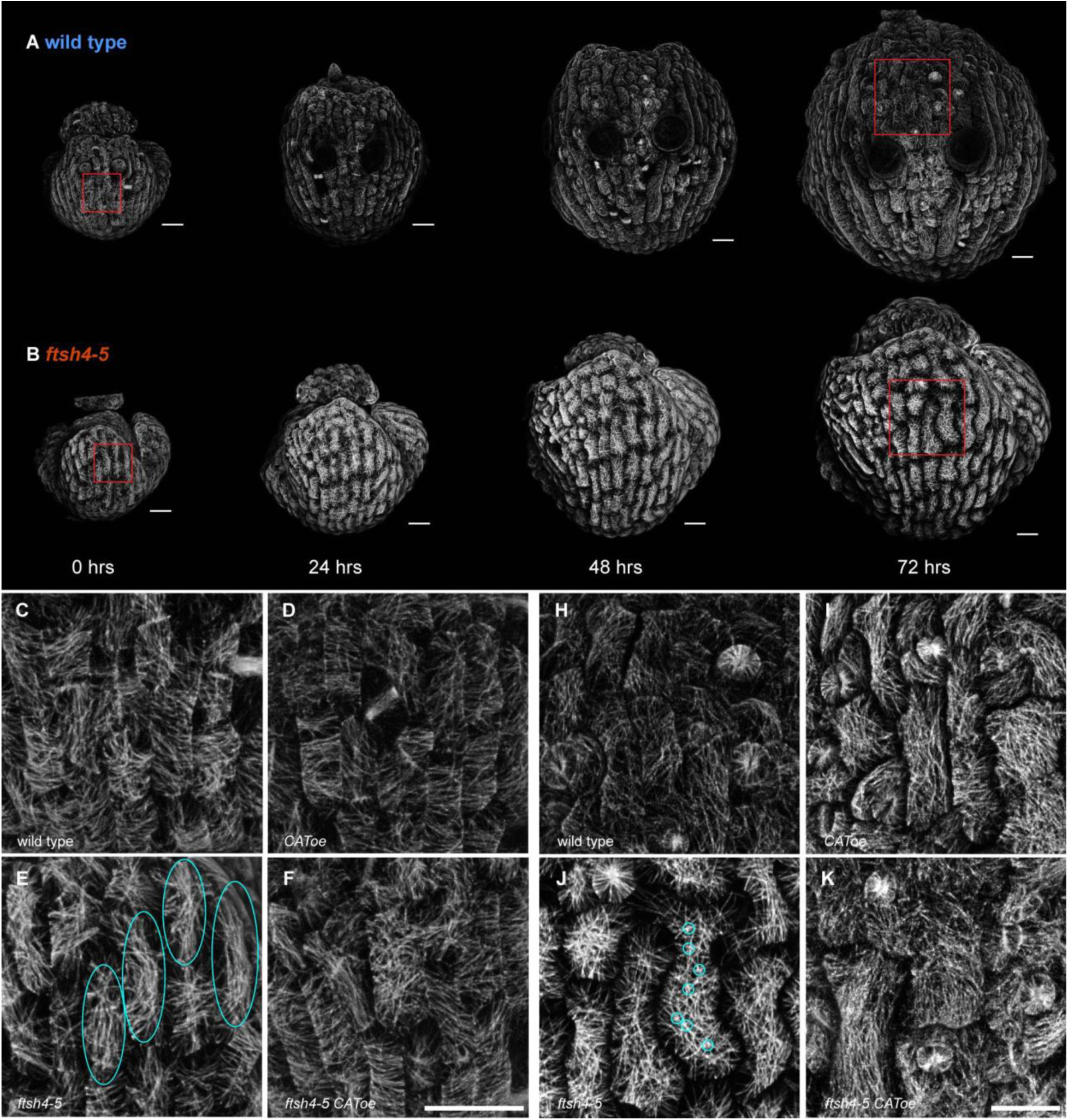
Microtubule arrangement is different in *ftsh4-5* and rescued in *ftsh4-5 CAT2oe*. A-B: Microtubule signal from the live time lapse imaging of sepal development. Representative replicates of (A) wild type, and (B) *ftsh4-5* from n=3 sepals. Scale bars are 20 μm. Red boxes indicate the location of the zoomed-in panels. C-F: Microtubule arrangement at the first day of imaging (developmental stage 5) for (C) wild type, (D) *CAT2oe*, (E) *ftsh4-5*, (F) *ftsh4-5 CAT2oe*. Panels are zoomed-in to show a few cells from the time lapse imaging. Representative images from n=3 replicates. Scale bars are 20 μm. Cyan ovals in (E) point out cells with longitudinal microtubules. H-K: Microtubule arrangement at the last day of imaging (developmental stage 7) for (H) wild type, (I) *CAT2oe*, (J) *ftsh4-5*, (K) *ftsh4-5 CAT2oe*. Panels are zoomed-in to show a few cells from the time lapse imaging. Representative images from n=3 sepals. Scale bars are 20 μm. Cyan circles in (J) point out the ‘star-shapes’ in one cell from many microtubules crossing over each other.

At the end of the time series (fourth time point, corresponding to floral stage 7), microtubules in wild type are isotropic (Fig 2H), as are microtubules in *CAT2oe* sepals (Fig 2I). The microtubule arrangement in *ftsh4-5* appears different in some patches of cells (Fig 2J). Although the microtubules are isotropic, there are many locations where the microtubule signal crosses over from many angles, so that its signal looks star-shaped. We describe this microtubule arrangement as ‘crisscrossed;’ it is an extreme degree of isotropy. The microtubule arrangement is rescued in *ftsh4-5 CAT2oe* (Fig 2K) and usually shows levels of isotropy like those in wild type. Therefore, microtubule arrangements are similar between genotypes with normal growth heterogeneity at the same developmental stage, whereas *ftsh4-5* microtubules have distinctly different microtubule arrangements at each developmental stage.

To quantify differences in microtubule arrangement between genotypes at all time points, we used machine learning to classify the microtubule signal as organized (anisotropic) or crisscrossed (Fig 3). The algorithm was trained to classify microtubules that cross over each other many times as crisscrossed, and instances of this can occur in all genotypes. The organized microtubules usually had a transverse orientation, which is labeled in blue in the example (Fig 3, top right). The crisscrossed microtubules in *ftsh4-5* are correctly labeled as crisscrossed in red (Fig 3, bottom right). We found that *ftsh4-5* on average had more crisscrossed microtubules at each time interval compared to other genotypes. *CAT2o*e had the most organized microtubule signal, and *ftsh4-5 CAT2oe* is a partial rescue with scores between wild type and *ftsh4-5* (Fig 3). The wild type, *CAT2oe*, and *ftsh4-5 CAT2oe* cells also have plenty of regions with isotropic microtubules that are sometimes also classified as crisscrossed. However, it is more common in these genotypes to have cells with both organized and crisscrossed microtubules (Fig 3, top left) whereas in *ftsh4-5* it is common to have entire cells with crisscrossed microtubules. At the 24 hr and 48 hr time points, 2 of 3 *ftsh4-5* replicates have crisscrossed microtubules, and the remaining replicate has longitudinal microtubules at the second day of imaging and then crisscrossed microtubules at the third day of imaging (Fig 2B, Fig S2F-G). Longitudinal microtubules in *ftsh4-5* were often classified as crisscrossed (Fig 3, bottom left), because they cross over each other locally, despite the cell-scale longitudinal organization. This makes sense considering that cells with longitudinal microtubules transition to having crisscrossed microtubules. All other genotypes have microtubules that progressively become more isotropic each day, except for one *ftsh4-5 CAT2* replicate which has a mild crisscrossed microtubule phenotype (Fig 2A, Fig S2A-E, H-J). Therefore, machine learning confirms that *ftsh4-5* has an abnormal crisscrossed microtubule arrangement, which is rescued when ROS is lowered.

**Figure 3:**
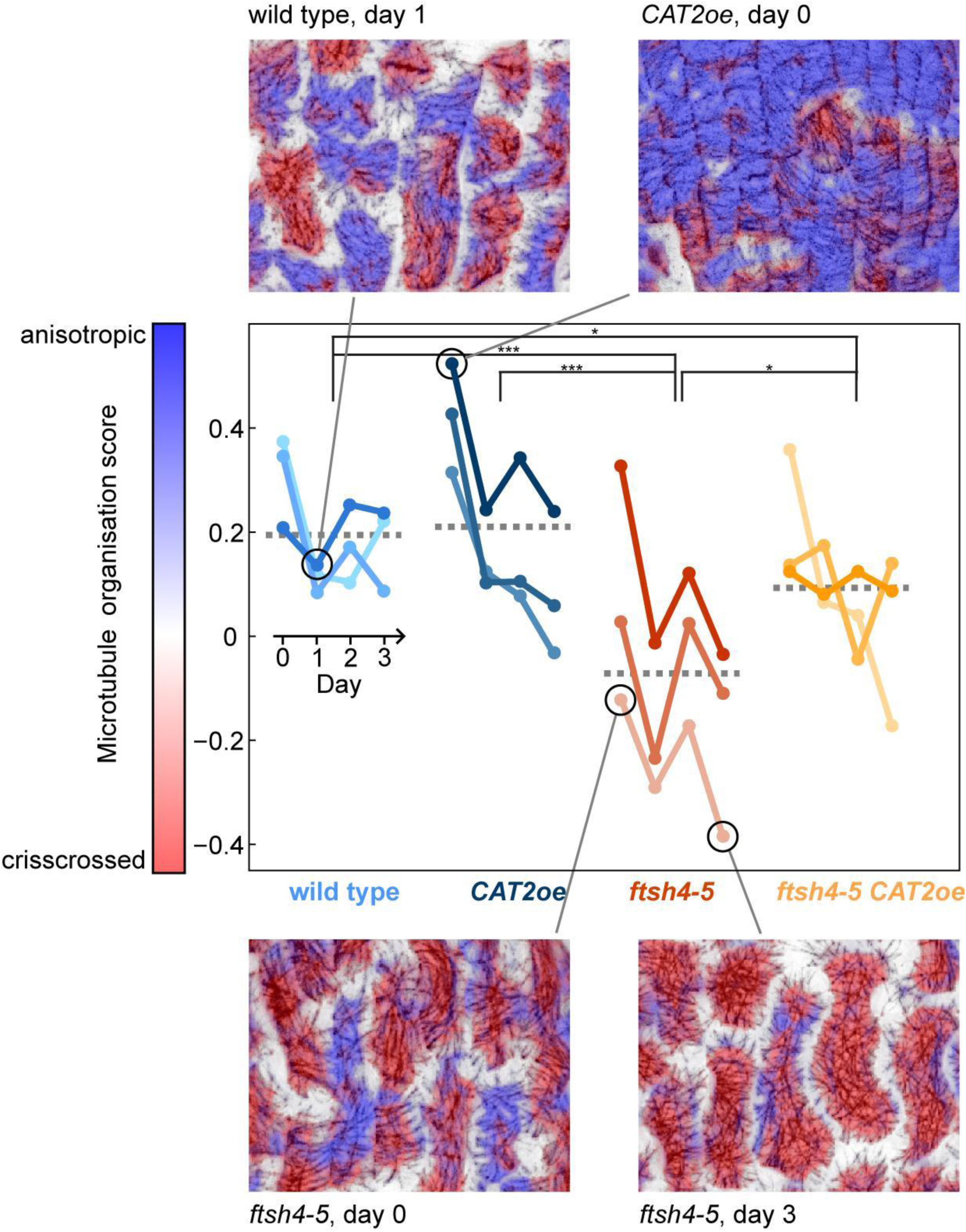
Microtubules in *ftsh4-5* are more crisscrossed. Machine learning was trained to classify microtubules as organized or crisscrossed on a subcellular scale using the microtubule signal from the live imaging. Then proportion of pixels classified as organized (represented in blue) or crisscrossed (represented) in each replicate and time interval was quantified. Pixels that are white were excluded from the analysis because they often contain distorted signal from microtubules on the anticlinal walls. * means p value <0.05, ** means p value <0.01, ***, and means p value <0.001.

To determine whether the elevated ROS in *ftsh4-5* is sufficient to cause crisscrossed microtubules, we elevated ROS in wild-type plants using the CATALASE inhibitor 3-AT (3-amino-1,2,4-triazole). Plants with the microtubule marker in the wild-type background were treated for 5 days, which is effective in recapitulating the variable organ size and shape phenotype (Fig 4A-B) and elevating hydrogen peroxide compared to the mock treatment (Fig 4C-F, Fig S3). The microtubules in mock-treated and 3-AT plants were also imaged after 5 days of treatment (Fig 4G-H, Fig S3I-J) Plants treated with 3-AT had microtubules that are brighter and crisscrossed (Fig 4H,I Fig S3J), in a manner that is strikingly similar to the microtubule arrangement in *ftsh4-5*. The microtubules in the mock treatment were isotropic with a slight bias in the transverse direction (Fig 4G,I Fig S3I) similar to untreated wild type. The recapitulation of crisscrossed microtubule arrangement by increasing ROS with 3-AT and the rescue of microtubule arrangement by decreasing ROS with *CAToe ftsh4-5* indicates that ROS causes the microtubule phenotype in *ftsh4-5* and is sufficient to cause the crisscrossed microtubules.

**Figure 4:**
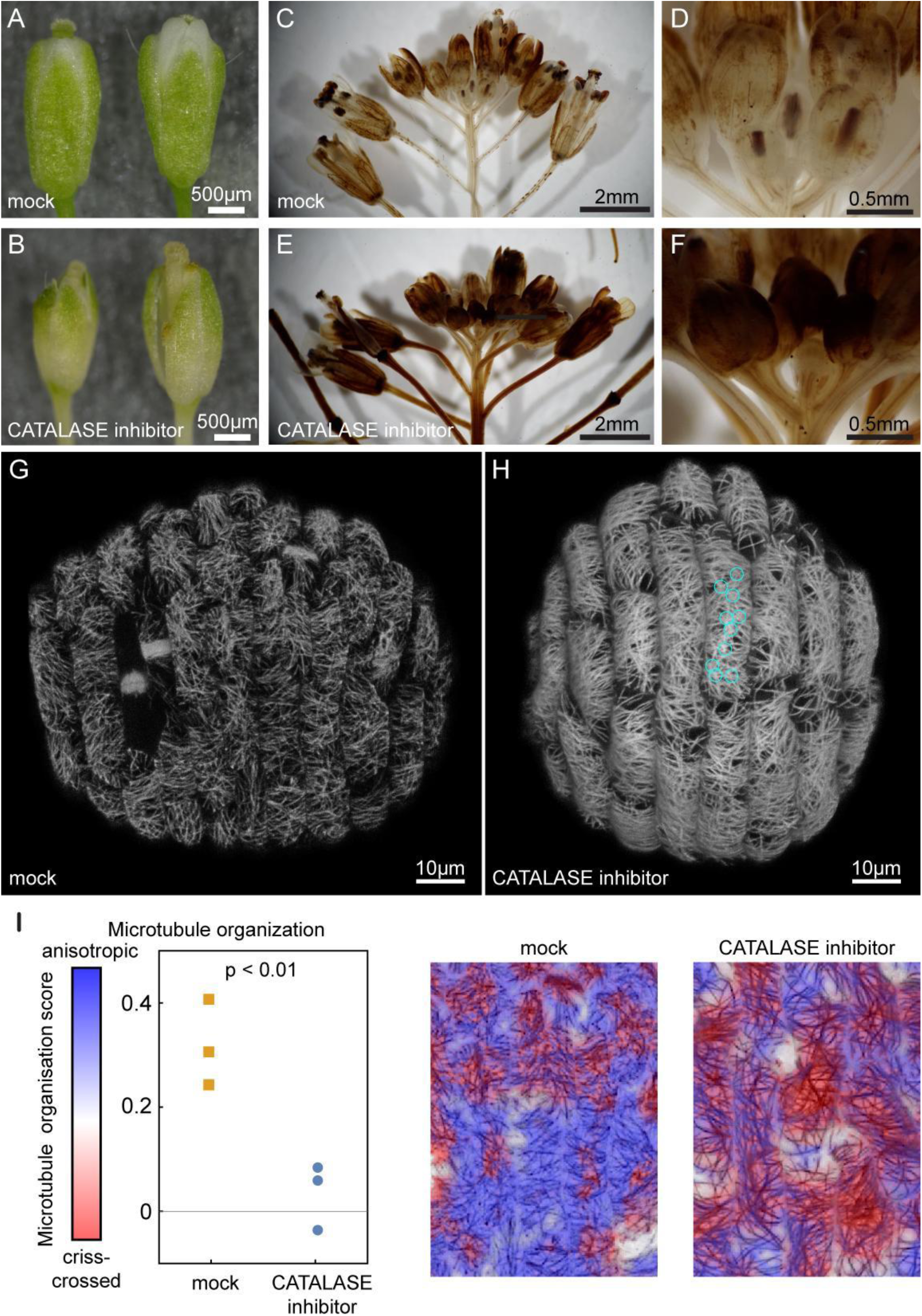
ROS is sufficient to lead to crisscrossed microtubules. A-B: Flowers from mocked-treated (A) and 3-AT treated (B) inflorescences. 3-AT treatment causes variability in organ size and shape. C-F: Inflorescences are stained for hydrogen peroxide with DAB. (C) Whole mock treated inflorescences have hydrogen peroxide in the sepals of flowers in later development and mature flowers as seen previously. (D) Magnification of the youngest flowers shows that hydrogen peroxide is low. (E) Whole 3-AT treated inflorescences have elevated hydrogen peroxide levels throughout, and (F) magnification of the youngest flowers shows that hydrogen peroxide is elevated in the youngest flowers as well. Representative images from n=3 replicates. G-H: Microtubules in (G) mock-treated and (H) CATALASE-inhibitor treated sepals. Cyan circles point out star shapes from crisscrossed microtubules in one cell. Representative images from n=3 replicates. I: Plot of the proportion of microtubules classified as crisscrossed (red pixels) or anisotropic (blue pixels) in the mock vs CATALASE-inhibitor treated sepals. Representative images show that red pixels are classified as crisscrossed and blue pixels are classified as anisotropic. White pixels were excluded from the analysis because they often contain distorted signal from microtubules on the anticlinal walls.

### Microtubules are more dynamic in wild type than in *ftsh4-5*

Next we tested whether the different microtubule arrangements in wild type and *ftsh4-5* corresponded to a difference in microtubule dynamics. To test the stability of microtubules, we used a propyzamide treatment which prevents microtubule polymerization (Akashi et al., 1988). More stable microtubules persist longer before depolymerizing. Thus, comparing the amount of microtubule depolymerization in a given amount of time allows for comparison of microtubule stability. Microtubules in wild-type and *ftsh4-*5 sepals at stage 5-6 (corresponding to the second day of live imaging) were imaged before and after a 30 min propyzamide treatment (Fig 5A-H, Fig S4). This length of treatment was chosen because it was long enough for partial depolymerization of microtubules. Microtubules in wild type appeared isotropic before treatment (Fig 5A-B, Fig S4A), and after treatment most microtubules were depolymerized (Fig 5C-D, Fig S4B). In the cells observed, microtubules in *ftsh4-5* appeared isotropic before treatment (Fig 5E-F, Fig S4C). The treatment caused depolymerization, but in contrast to wild type, most cells still had some microtubules remaining (Fig 5G-H, Fig S4D). Since microtubules in *ftsh4-5* depolymerized less than in wild type, this suggests that *ftsh4-5* cortical microtubules are more stable. As a control, the microtubules in the preprophase bands depolymerized completely in both genotypes, indicating that this is not a difference in sensitivity to the treatment (Fig S4 E-H). The organization of the microtubules that remained after treatment also differed between wild type and *ftsh4-5*. In wild type, remaining microtubules were often anisotropic in the transverse direction (Fig 5D) even when microtubules were isotropic before treatment (Fig 5B), whereas the remaining microtubules in *ftsh4-5* were usually isotropic (Fig 5H) and had a similar organization before and after treatment (Fig 5F). When the arrangement of the microtubules was scored per cell after treatment, wild type had more cells with fully depolymerized microtubules, and more cells that had anisotropic microtubules remaining compared to *ftsh4-5* (Fig 5I). These results suggest that there is a difference in microtubule stability, and a difference in the orientation of the most stable microtubules between wild type and *ftsh4-5*.

**Figure 5:**
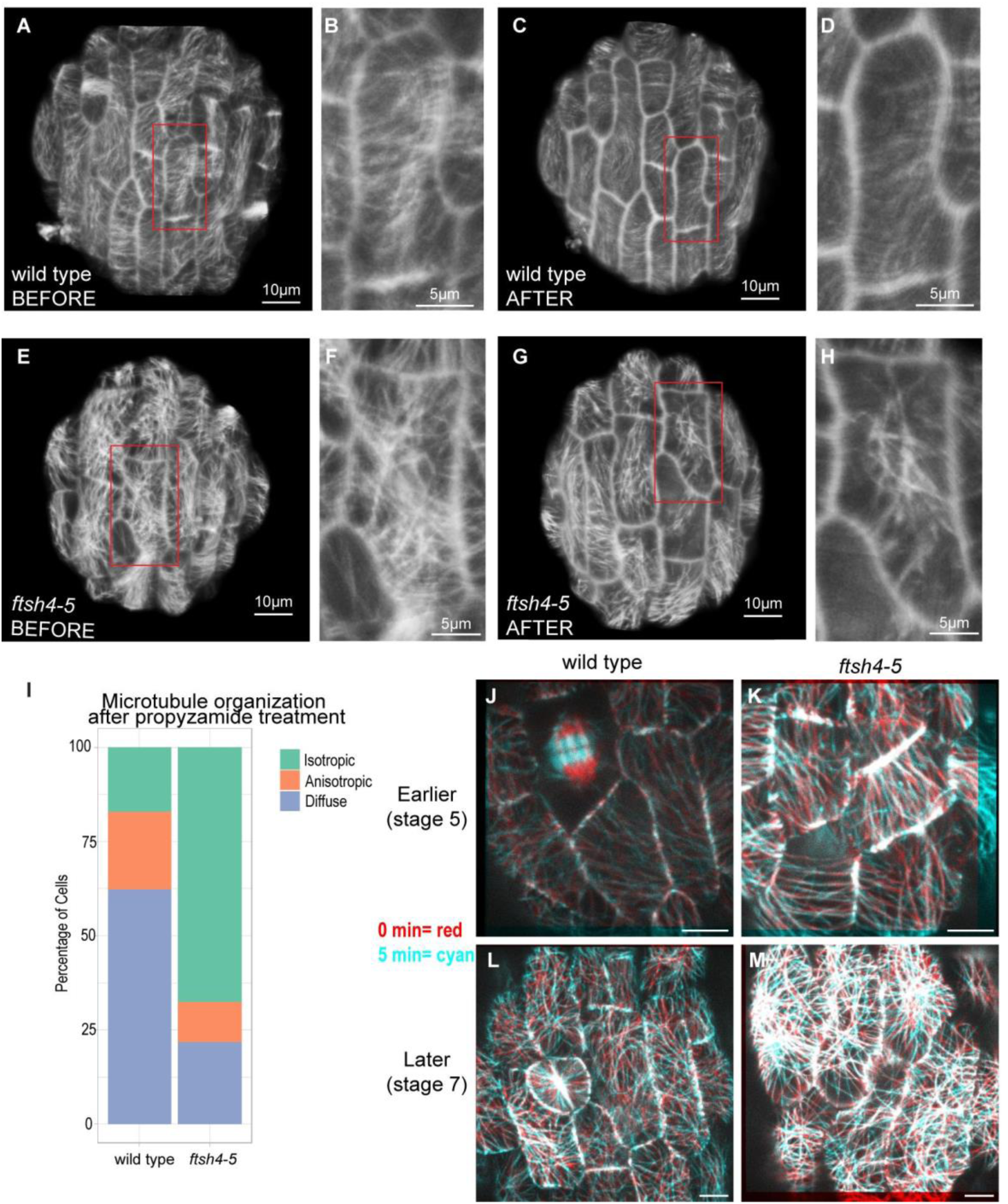
Crisscrossed microtubules in *ftsh4-5* are more stable. A-I: Propyzamide treatment to compare microtubule stability. (A) Wild-type sepal cells before treatment, and the red box marks the location of the magnification in (B). (C) The same wild-type sepal cells after 30 min propyzamide treatment, and the red box marks the location of the magnification in (D). (E) *ftsh4-5* sepal cells before treatment, and the red box marks the location of the magnification in (F). (G) The same *ftsh4-5* sepal cells after 30 min propyzamide treatment, and the red box marks the location of the magnification in (H). (I) The arrangement of microtubules in each cell was scored after propyzamide as anisotropic, isotropic, or depolymerized polymerized (which appears as diffuse signal in the cell). Scores are displayed as percentage of total cells in a stack bar graph for wild type and *ftsh4-5*. J-M: Microtubules are imaged every 5 min and displayed as a merge of the 0 min time points in red and the 5 min time points in cyan. Separated colors indicate changes in microtubule orientations. Red and Cyan merge into white which indicates microtubules that are present at both time points. Microtubules are imaged in wild type (J) and *ftsh4-5* (K) at the developmental stage corresponding to the first day of the live imaging time series (stage 5). Representative images of n=3 replicates. Scale bar is 4μm. Microtubules are imaged in wild type (L) and *ftsh4-5* (M) at the developmental stage corresponding to the last day of the live imaging time series (stage 7). Representative images of n=3 replicates. Scale bar is 4μm.

To examine microtubule dynamics more directly, we imaged microtubules once every 5 minutes. In both wild-type and *ftsh4-5* sepals of at the stage corresponding to 0 hr time point in the live imaging series (stage 5) there were many changes in individual microtubules in each 5 min interval (Fig 5J-K, Video 1). At this developmental stage, *ftsh4-5* microtubules are often longitudinal (Fig 2E). In some cells with longitudinal microtubules, the microtubules reorganize to be more crisscrossed for one or a few frames, and then again become longitudinal (Video 1). This suggests that there is not a unidirectional transition from longitudinal to crisscrossed microtubules in *ftsh4-5*. Our analysis suggests that microtubules in *ftsh4-5* and wild type are dynamic at this stage of sepal development, with many microtubules changing within 5 min time intervals.

In the developmental stage that corresponds to the last day of the live imaging series (stage 7), wild type microtubules are isotropic and still have many changes over each 5 min interval (Fig 5L, Video 2). In *ftsh4-5*, while there are still changes in microtubules over each 5 min, crisscrossed microtubules persist longer than the other microtubules (Video 2, Fig 5M). This result indicates that crisscrossed microtubules in *ftsh4-5* have increased stability. It is likely that these less dynamic crisscrossed microtubules that persist longer in *ftsh4-5* are the same stable isotropic microtubules that remained in *ftsh4-5* after propyzamide treatment. Also, it should be noted that despite substantial dynamics in individual microtubules, the cell-scale general orientation of microtubules did not change over the 65 min total imaged in either development stage or genotype, suggesting that 24 hr time intervals are still informative of average cell-scale microtubule direction. Altogether the results of these experiments suggest that wild type microtubules continue to be dynamic during sepal development, whereas crisscrossed microtubules in *ftsh4-5* have enhanced stability.

### Cells with temporally correlated growth have crisscrossed microtubules

We next tested whether the differences in the arrangement and stability between wild type microtubules and *ftsh4-5* crisscrossed microtubules correlated with differences in cell growth in our live imaging time series. We compared microtubule arrangements in cells with normal growth in wild type (Fig 6A), cells in slow-growing and temporally correlated patches in *ftsh4-5* (Fig 6B), and cells in normal-growing patches in *ftsh4-5* (Fig 6C). Cells in slow-growing patches in *ftsh4-5* have longitudinal microtubules at earlier time points and crisscrossed microtubules at later time points (Fig 6B). However, microtubules in normal-growth patches in *ftsh4-5* looked similar to wild type (Fig 6C). Since *ftsh4-5* has a range of severity in its phenotype, we also looked at the correlation between the organ-scale microtubule organization and growth heterogeneity in individual replicates. We tested whether the microtubule organization is associated with the previous quantifications of temporal correlation in growth and patchiness in growth accumulation. We find that replicates with more crisscrossed microtubules have more temporal correlation in growth fluctuations (Fig 6D) and patchier accumulation of growth (Fig 6E). This indicates that the abnormal microtubule arrangement in some *ftsh4-5* cells is associated with the slow, temporally correlated growth patches.

**Figure 6:**
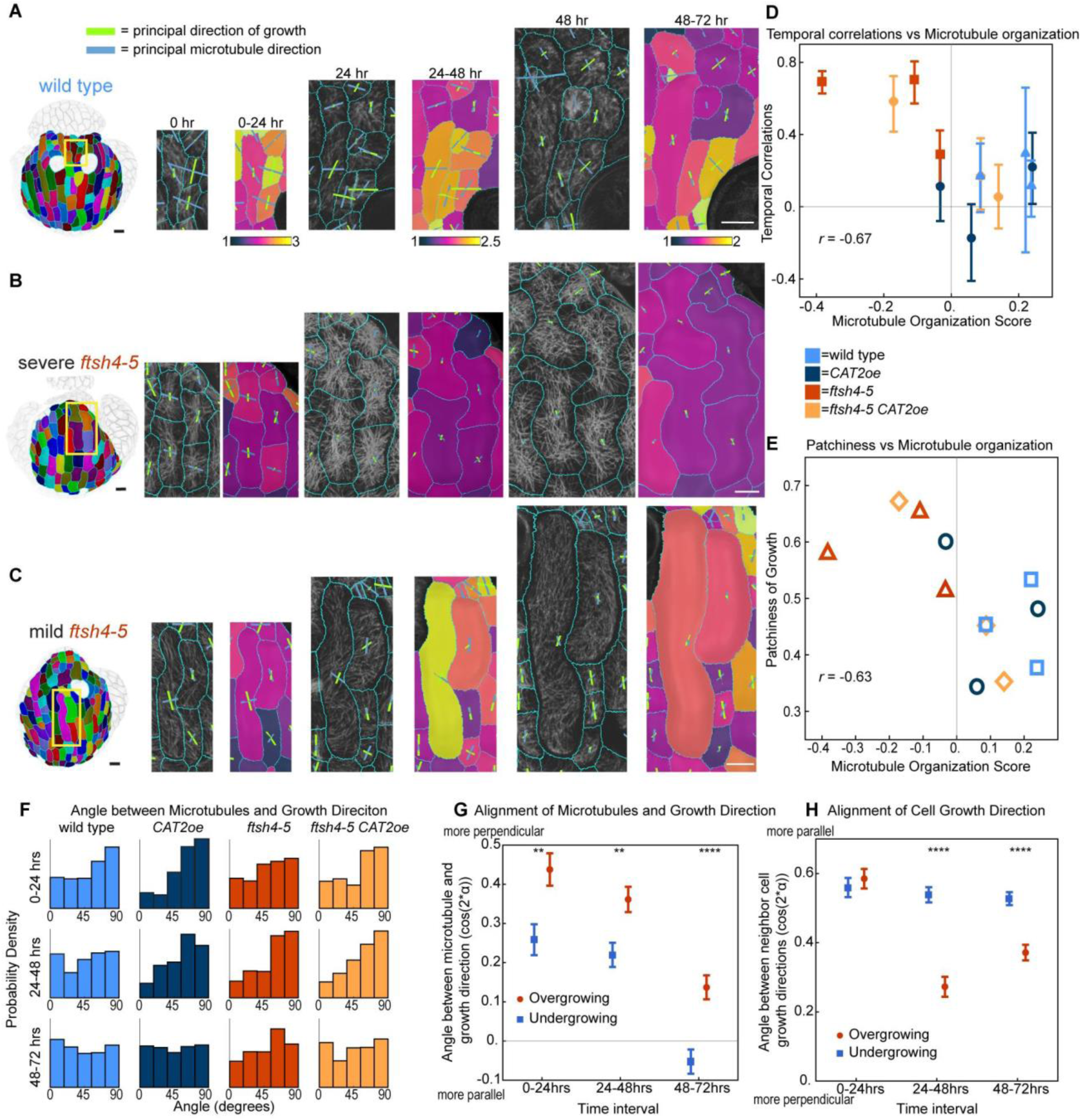
Cells with decreased growth heterogeneity have crisscrossed microtubules, and growth direction is biased towards perpendicular to microtubule direction. A-C: Microtubule direction, growth rate, growth direction can be obtained from the same live imaging data set. Scale bars are 10μm. (A) Wild type sepal cells with heterogeneous growth rate have transverse or isotropic microtubules. (B) Cells with decreased growth heterogeneity in *ftsh4-5* have microtubules that are longitudinal and then crisscrossed. (C) Cells heterogeneous growth in *ftsh4-5* have microtubules that look more similar to those in wild type. (D) The score for temporal correlation in growth fluctuations from Fig 1N is plotted against microtubule organization score from Fig 3 for individual replicates. Correlation coefficient is -0.67. (E) The score for patchiness in growth fluctuations from Fig 1O is plotted against microtubule organization score from Fig 3 for individual replicates. Correlation coefficient is -0.63. (F) Weighted histograms of the angle between microtubule direction per cell and cell principal direction of growth plotted for each genotype and time interval. Histograms are also weighted by microtubule anisotropy and growth direction anisotropy by (max - min)/(max + min) for both values and using the product of these values as weights. (G) Plot of correlation between microtubule direction and growth direction for cells overgrowing relative to their neighbors and undergrowing relative to their neighbors. Data from all genotypes is pooled. Angle is represented as –cos(2*alpha). This is –1 if alpha = 0, i.e., microtubule and growth direction are parallel for all cells and 1 if alpha = pi/2, i.e. microtubule and growth direction are perpendicular for all cells. (H) Plot of correlation between neighboring cell growth directions for cells overgrowing relative to their neighbors and undergrowing relative to their neighbors. Data from all genotypes is pooled. Angle is represented as cos(2*alpha). This is -1 if alpha = pi/2, i.e. neighboring growth directions are perpendicular for all cells and 1 if alpha = 0, i.e. neighboring growth directions are parallel for all cells. ** means p value <0.01 and **** means p value<0.0001.

### Growth direction is biased towards perpendicular to microtubule direction

Cell walls are expected to deform perpendicular to cellulose microfibril direction (Coen and Cosgrove, 2023), and therefore it is often assumed that cell growth direction is perpendicular to microtubule direction. However, wild-type sepal cells often have isotropic microtubules despite organ elongation in the proximal-distal direction. Since there is heterogeneity in growth direction between cells (Burda et al., 2024; Hong et al., 2016), we tested whether cell growth direction and microtubule orientation were correlated on a cell-scale. The analysis was also weighted by the anisotropy of microtubule direction and growth direction, since it is expected that microtubule anisotropy would cause growth anisotropy. We found that the angle between principal direction of cell growth over 24 hours and principal microtubule orientation at the start of the time interval is biased towards 90 degrees in many of the genotypes and time intervals (Fig 6F). Growth during the 0 to 24hr time interval in wild type, *CAT2oe*, and *ftsh4-5 CAT2oe* have growth directions that are the most biased towards perpendicular to microtubules (Fig 6F). To further resolve the relationship between microtubule and growth direction, we tested whether it was affected by growth rate. We find that the angle between microtubule and growth direction is significantly closer to perpendicular in cells that are overgrowing than cells that are undergrowing relative to their neighbors (Fig 6G). This result suggests that increased growth rate may cause microtubules to have greater influence on cell growth direction. We then hypothesized that growth direction is also influenced by the growth directions of neighbor cells, as cells cannot move past each other due to the cell walls.. To test this, we measured the angle between growth directions of neighboring cells and found that cells that are overgrowing have growth directions that are less aligned with those of the neighboring cells (Fig 6H). This means that neighboring cells can influence growth direction, and that faster growth overcomes some of this influence. Together, these results suggest that cell growth direction is influenced by both microtubule direction in a cell and neighboring cell growth directions. Faster growth causes growth direction to be influenced more by its own microtubule direction rather than the neighboring cells.

### Depolymerization of microtubules is not sufficient to restore normal cell growth fluctuations

Since cells with stable microtubules also display increased correlation of growth fluctuations, we tested whether removing microtubules could decrease correlations in growth by depolymerizing microtubules with oryzalin. We time-lapse live-imaged wild type and *ftsh4-5* sepal development once every 24 hours for 4 days, as was done for the previous live-imaging series (Fig 7A-D, Fig S5A-J). The oryzalin treatment began one day before the start of imaging, so microtubules were partially depolymerized on the first day of imaging and fully depolymerized on the second day of imaging (Fig S6). We quantified the temporal correlations in cell growth fluctuations and found that oryzalin treatment does not affect the temporal correlations in growth in neither wild type nor *ftsh4-5* (Fig 7E). The mock treatment had little effect on wild type (Fig 7A, Fig S5A-B) or *ftsh4-5* sepal growth (Fig 7B, Fig S5C-D). Oryzalin treatment (Fig 7C-D, Fig S5E-J) caused the cells to become more isotropic in shape as previously reported (Hamant et al., 2008; Zhu et al., 2020). This presumably occurs because cellulose microfibril orientation is disrupted so that cellulose cannot constrain growth, and so cells expand isotropically. This demonstrates that growth heterogeneity is not restored immediately when microtubules are depolymerized.

**Figure 7:**
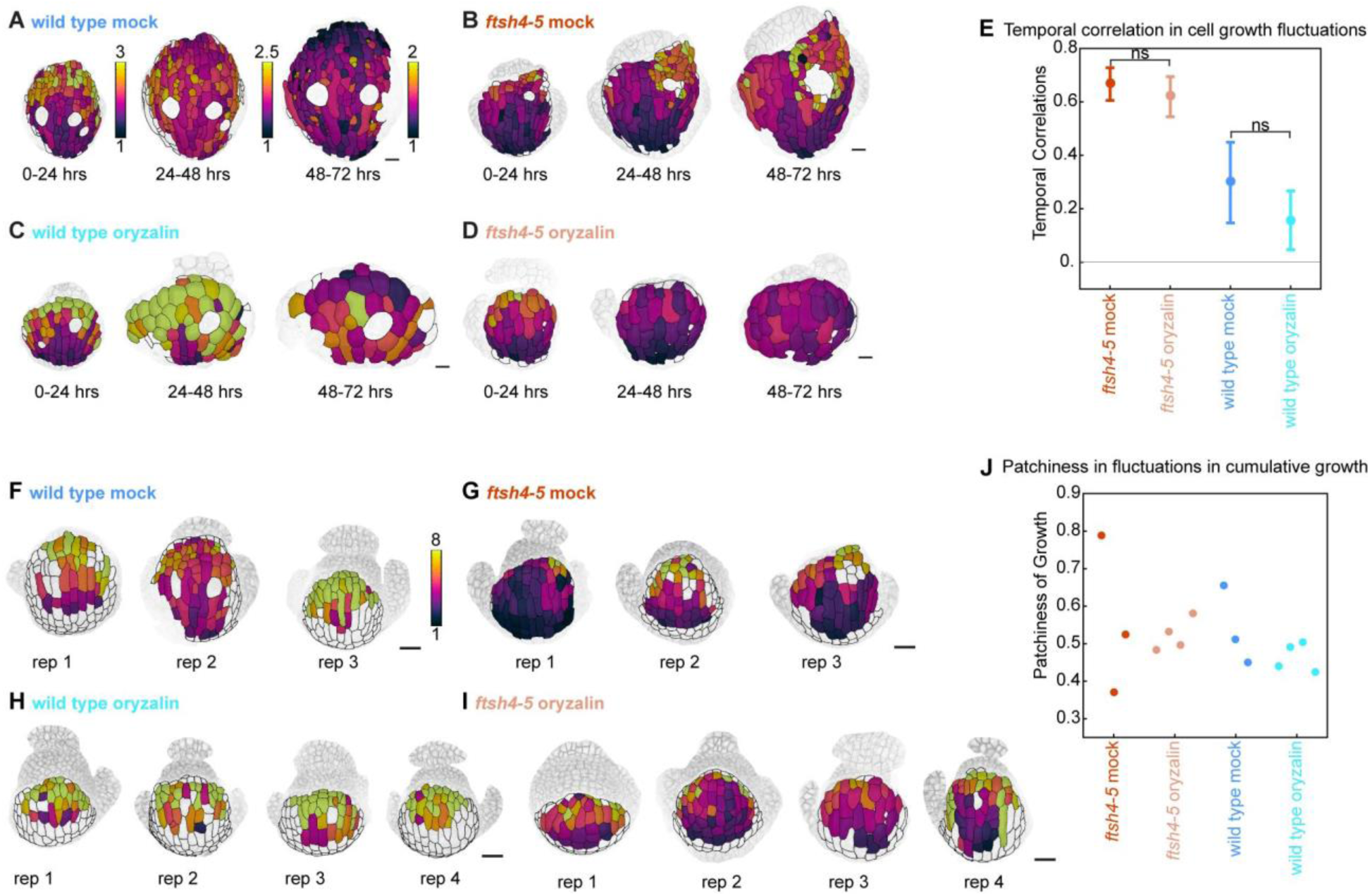
Depolymerizing the microtubules is insufficient to restore growth heterogeneity. A-D: Cell area growth heat maps of sepal development with an oryzalin treatment. Sepals were imaged once every 24 hours for 4 days. Area growth is represented as a ratio and projected onto the later time point. Representative time series for (A) wild type mock treatment, (B) wild type oryzalin treatment, (C) *ftsh4-5* mock treatment, (D) *ftsh4-5* oryzalin treatment. n=3 for mock treatments and n=4 for oryzalin treatments to account for fewer cells from decreased division. (E) The fluctuations in growth rate around the average growth for each time point and replicate are calculated, and then the temporal correlation in fluctuations over subsequent 24 hr intervals is calculated. ns means nonsignificant and differences between all other groups is significant. F-I: Heat maps of cumulative cell area growth over 3 days for all replicates of (F) wild type mock treatment, (G) *ftsh4-5* mock treatment, (H) wild type oryzalin treatment, and (I) *ftsh4-5* oryzalin treatment. (J) Patchiness of growth over 3-days is calculated by the standard deviation of the fluctuations after averaging the fluctuation over the immediate neighboring cells.

To assess whether microtubules affect spatiotemporal averaging of growth rate fluctuations, we quantified the patchiness of the cumulative growth. Mock treatment does not affect cumulative growth, which is evenly distributed in wild type (Fig 7F), and unevenly distributed in *ftsh4-5* (Fig 7G). Oryzalin did not have an obvious effect on patchiness of growth (Fig 7J). In oryzalin-treated wild type, cumulative growth is also evenly distributed (Fig 7H). In oryzalin-treated *ftsh4-5* (Fig 7I), cumulative growth is very low in 3 of 4 replicates because slow-growth patches encompass most of the sepal, and patchy in 1 of 4 replicates. This indicates that depolymerizing microtubules cannot rescue the loss of spatiotemporal averaging in *ftsh4-5*.

## Discussion

Cell growth heterogeneity occurs during morphogenesis and needs to be averaged in order for development to be robust. The *ftsh4-5* mutant has a loss of robust development due to temporally correlated growth fluctuations that inhibit spatiotemporal averaging (Burda et al., 2024; Hong et al., 2016). The loss of developmental robustness is due to elevated ROS (Hong et al., 2016). Various other factors have been linked to growth rate heterogeneity, including microtubules which indirectly affect the mechanical properties of the cell wall. We find that elevated ROS promotes temporal correlations in growth fluctuations which also inhibits spatiotemporal averaging of growth fluctuations. Elevated ROS are also sufficient to cause microtubules to become ‘crisscrossed,’ and crisscrossed microtubules in *ftsh4-5* have increased stability. Cells with crisscrossed microtubules also exhibit temporally correlated growth fluctuations. However, depolymerizing microtubules was insufficient to decrease temporal correlations in growth fluctuations, likely due to existing cellulose in the cell wall before treatment.

### Stable microtubules may have increased bundling

The increased stability and thicker appearance of crisscrossed microtubules in *ftsh4-5* suggests that they are bundled. In sepals, the curvature of the tissue limits the resolution of microtubule imaging due to the curvature of microtubules in the z plane and because the 3D structure of a flower bud limits the options for mounting samples for imaging. Therefore, we were unable to observe microtubule dynamics at a resolution that would have allowed us to observe bundling of individual microtubules directly.

However, aspects of *ftsh4-5* microtubule dynamics are similar to those observed for bundling, such as thicker appearance and increased stability (Van Damme et al., 2004). Crisscrossed microtubules in *ftsh4-5* cross over each other at large angles so that many cross over each other to form a star shape. Typically, in wild type, microtubules that cross over at large angles lead to severing whereas crossing over at shallow angles lead to bundling, and this leads to organization (Dixit and Cyr, 2004). MAP65-1 mediates microtubule bundling, and this type of bundling is protective against severing by KATANIN1 (Burkart and Dixit, 2019). Changing the rate of severing or the stability of microtubules also changes microtubule organization (Dixit and Cyr, 2004; Zhang et al., 2013). Therefore, it is possible that MAP65-1 activity, or the activity of a similar protein, could create the crisscrossed pattern in *ftsh4-5* by inhibiting severing of microtubules. There is also evidence, in maize, that ROS increases the expression of MAP65-1 (Zhu et al., 2013). Thus, we hypothesize that increased stability of *ftsh4-5* microtubules is likely due to increased bundling, and that bundling may lead to the disorganized crisscrossed microtubule pattern. In the future, it will be interesting to test whether MAP65-1 or other MAPs contribute to crisscrossed microtubules *ftsh4-5*.

### Complex relationship between growth direction and microtubule orientation

It is expected that cellulose microfibrils are assembled parallel to cortical microtubules, and that cell growth is perpendicular to cellulose microfibrils. We found that the angle between cell growth direction and average microtubule direction is biased towards perpendicular. We also found that microtubule direction and growth direction are closer to perpendicular in cells that are growing faster than their neighbors. The growth directions of cells growing slower than their neighbors is more influenced by the growth direction of the neighboring cells. Surrounding cells constrain growth and affect the relationship between microtubule and growth direction. However, the relationship is not completely resolved as there are faster growing cells with growth that is not perfectly perpendicular to microtubule direction. There is the possibility that the time scale on which microtubules affect the growth direction is shorter than 24 hours and we did not capture this effect, or that previously synthesized cell wall components have an influence on growth direction which is not accounted for here. Previously synthesized cellulose also has an effect on growth direction. The *csi1* mutant, in which cellulose synthase complexes are not tethered to the microtubules, has more anisotropic new cellulose, but similar anisotropy in total cellulose compared to wild type, and has similar growth anisotropy as wild type (Mollier et al., 2023). We also imaged only one outer wall of each cell studied. The inner cell wall could contribute to the direction of cell growth. Further, the side (anticlinal) walls have been shown to be important for anisotropic growth and flattening of the organ (Zhu et al., 2020), and they are also not accounted for here. Supracellular tensile stress also likely affects microtubule organization (Burian et al., 2013; Hervieux et al., 2016; Uyttewaal et al., 2012), and microtubules align better with the predicted direction of tensile stress than the cell growth direction in the boundary region of the shoot apical meristem (Burian et al., 2013).

### Relationship between microtubule dynamics and growth fluctuations

Here we found that depolymerizing microtubules was insufficient to restore growth rate heterogeneity in the subsequent days. However, if cellulose microfibrils affect growth for a few days or longer, then the previous microtubule organization and cellulose microfibril organization would still affect the cells with no currently polymerized microtubules. Previously synthesized cellulose can also guide the cellulose synthase complexes in the absence of microtubules (Paradez et al., 2006). This may explain why depolymerizing the microtubules did not affect correlations in growth fluctuations, yet it was previously reported that decreased microtubule dynamics in *katanin* does decrease heterogeneity in cell growth rates (Uyttewaal et al., 2012).

It is also possible that microtubule dynamics are necessary for creating growth heterogeneity, and depolymerizing microtubules further decreases microtubule dynamics. There is evidence that microtubule dynamics can generate growth heterogeneity; increased growth of a differentiating trichome cell causes increased tension, increased microtubule anisotropy, and decreased cell growth rate in the surrounding cells (Hervieux et al., 2017). Microtubule dynamics promote anisotropic arrangements (Dixit and Cyr, 2004) and the ability to change the orientation of microtubules over time (Uyttewaal et al., 2012). Perhaps changes in microtubule anisotropy and direction creates changes in cell wall properties over time and space, leading to fluctuations in cell growth rates. On the other hand, more stable microtubules may not generate changes in cell wall properties over time and space, leading to increased correlation in fluctuations in cell growth rates. In this scenario, microtubule response to tension created by growth fluctuations would propagate growth fluctuations, leading to spatiotemporal averaging of fluctuations and robust development of organ size and shape. Perhaps depolymerizing the microtubules in *ftsh4-5* did not cause any further changes in the anisotropy of cell wall properties, which assumably would have been necessary to change the correlations in growth fluctuations.

### Microtubule organization could be a consequence of tensile stress in the cell

Cells in *ftsh4-5* sometimes become lobed (indented in places to create a more complex shape) (Sapala et al., 2018). Usually, these cells are in the slow-growing patches that have decreased temporal growth heterogeneity. Leaf epidermal cells are also lobed to the extent that they are described as ‘puzzle-shaped.’ The development of lobes mediates tensile stress by transferring it from the center of the cell to the indented regions of the cell (Sampathkumar et al., 2014; Sapala et al., 2018). The indented regions have increased microtubule anisotropy, which is likely because microtubules align parallel to cell shape-derived stress (Sampathkumar et al., 2014). Therefore, it is possible that microtubules in sepal cells are influenced by stress created by cell geometry, especially in lobed *ftsh4-5* cells.

### ROS in sepal development

The elevated ROS levels in *ftsh4-5* could be disrupting a normal role of ROS in development. Although ROS is toxic at high levels, normal levels of ROS are important for redox biology and many cellular functions, and thus ROS levels are tightly regulated (Mittler, 2017). ROS localization has a spatial pattern in Arabidopsis sepals, in which ROS first appears at the tip of sepals and then progresses towards the base (Hong et al., 2016). This suggests that ROS also has a normal, regulated role in sepal development or maturation. ROS signaling is not the only mechanism by which metabolism and growth are linked. Nutrient levels and sensing also affect growth (Boulan and Léopold, 2021). The TOR complex is involved in sensing energy, and in Arabidopsis, decreased TOR activity affects protein translation and results in variable localization of auxin and cytokinin, which disrupts the timing and localization of sepal initiation (Kong et al., 2024). Thus, cell functions that occur throughout the life cycle can also have a role in growth heterogeneity and developmental robustness.

## Methods

### Plant material

The *Arabidopsis thaliana* accession Col-0 plants are used as wild type and all mutants are in Col-0 background as well. Isolation of the *ftsh4-5* mutant is described in Hong et al 2016. The membrane marker is the transgene *pUBQ10::mCherry-RCI2A*. The microtubule marker is the transgene *pUBQ10::GFP-MBD* where MBD is the microtubule binding domain of MAP4, and the transgene came from the lab of Olivier Hamant. The transgenes were crossed into *ftsh4-5*. The original *p35S::CATALASE2* plants from Hong et al 2016 were silencing expression of the transgene when crossed into the plants with microtubule and membrane markers. Thus, we did an LR reaction (LR clonaseII) with *CAT2* inserted into pDONR201 with pK7WG2 to make a new Kanamycin resistant *p35S::CAT2* (also named pBB7). We transformed *p35S::CAT2* into wild-type and *ftsh4-5* plants with both the membrane and microtubule markers. Individual T1s were used for experiments.

### Genotyping, plant selection and qPCR

Genotyping of *ftsh4-5* was done as described in Burda et al 2024 and Hong et al 2016. Membrane and microtubules marker plants were selected by screening for fluorescence. *p35S::CATALASE2* was selected by germination on media with kanamycin. The media contained of 2.2g/L Murashige and Skoog, 0.5g/L MES, 5g/L sucrose, the pH was brought to 5.7 with KOH, then 10g/L phytoagar was added, and after autoclaving kanamycin was added to the concentration of 50μg/ml. Leaf tissue was collected from individual T1 plants used for imaging and ROS stains, and then mRNA was extracted using the RNeasy Plant Mini Kit using the RLT buffer and beta-mercaptoethanol. Then cDNA was synthesized from the mRNA by doing a DNase treatment (NEB DNase1(RNase-free) catalog number M0303L) and then forward strand synthesis (Invitrogen Superscript II Reverse Transcriptase catalog number 18064014), and then used for qPCR. Three separate primer pairs targeting *CATALASE2* were used. Primer pair one is: 5’ GCACAGGGACGAGGAGGTTA 3’ and 5’ GCAGGCGGAGTTGGATACTT 3’. Primer pair two is: 5’ TGGAAAACGTGAGAGGTGCAT 3’ and 5’ TGCGGATTTCATGCGTGATG 3’. Primer pair three is: 5’ TCTTCAACCTGTTGGACGTATG 3’ and 5’ ATAGGAGAAGACACGGGTTTGA 3’. Fold change of each replicate was normalized to the average expression level of the three wild type replicates. Genotyping of the T2s to confirm segregation of *p35S::CATALASE2* was done with the forward primer in the 35S promoter (5’ caaccacgtcttcaaagc 3’) and the reverse primer in the *CAT2* coding sequence (5’ GATAACGGTGGAGAACCG 3’).

### Images of phenotypes and ROS stains

Images of flower morphology and ROS stains were taken with an Excelis 4K camera mounted on a Zeiss Stemi 508 stereomicroscope. Inflorescences stained for ROS were submerged in the same solution used for bleaching chlorophyll from the samples (3:1:1 ratio of ethanol: acetic acid: glycerol), arranged to spread out the flowers, and had a coverslip placed on top.

### ROS stains

3,3’-diaminobenzidine (DAB) and nitroblue tetrazolium (NBT) were used to stain for hydrogen peroxide and superoxide, respectively. Staining solutions and the bleaching solution were made as described in Hong et al 2016. Inflorescences were stained with DAB for 5 hours and with NBT for 24 hours, including vacuum infiltration for about 25 min.

### Microscopy and image analysis

Inflorescences were dissected and mounted in apex culture media (Hamant et al., 2014). Media containing 2.3 g/L Murashige and Skoog, 1% sucrose and 0.1% MES was brought to a pH of 5.8 with KOH, and agarose was added to a concentration of 1.2%. After autoclaving, media was supplemented with vitamins (final concentration of 100 µg/ml myoinositol, 1 ng/ml nicotinic acid, 1 ng/ml pyridoxine hydrochloride, 1 ng/ml thiamine hydrochloride, 2 ng/ml glycine). Plants then grew in 16 h light/8 h dark conditions on the media. Plants were switched to new media every 2-3 days. Abaxial sepals were imaged because they face outwards, making them the most accessible for imaging.

For the 24 hr live time lapse imaging, plants with the transgenes *pUBQ10::GFP-MBD* and *pUBQ10::mCherry-RCI2A* were dissected two days before imaging. Images were taken with a Leica Stellaris 5 using a 25X water dipping objective with an NA of 0.95 (HC FLUOTAR L VISIR 25X/0.95 WATER). A 488 laser with a power of 0.7 and a gain of 75, detecting wavelength of 494-550, was used to image GFP-MBD. A 561 laser with a power of 0.7 and a gain of 100, detecting wavelength of 582-607, was used to image mCherry-RCI2A. Signal from anthocyanin was collected at wavelengths 647-656 with a gain of 100 in a third channel. Channels were imaged in the same track, with line averaging of 2 and a scanning speed of 400, and bidirectional scanning. Images were a format of 1024x1024 pixels and 8 bit. The zoom was adjusted to fit the sepal and ranged from 2.25-3 on the first day of imaging and 1.6-2 on the last day of imaging. The z-step was 0.2µm. Inflorescences were positioned at an angle before each image and imaged once every 24 hrs.

MorphoGraphX was used to analyze cell growth and cortical microtubules. To remove red fluorescence from anthocyanin which interfered with the mCherry-RCI2A signal, the anthocyanin signal was added to itself to be twice as bright, and then subtracted from the mCherry-RCI2A channel. Then a neural network (VijayanUNET) was run on the ‘cleaned’ mCherry-RCI2A channel to predict locations of the cell wall, which creates a stack with a brighter and more continuous surface. This new stack was then used with an edge detect signal threshold to detect the surface. This was used to create the mesh, and then the signal from the cleaned mCherry-RCI2A was projected onto the mesh and used for segmentation. Cell lineages were tracked across time points, and then cell area growth and principal directions of cell growth, and anisotropy of growth direction were calculated. The cell neighborhoods were also saved. The proximal-distal axis was created by selecting cells at the tip of the sepal, creating a distance heat map from the selected cells (and saving the heat map), and then generating a custom axis from the heat map directions, and smoothing the axis. The principal directions of cell growth were added to the attribute map, and then the angle between the custom axis and the principal directions of growth was calculated. Then the microtubule signal was projected onto the mesh, cells with blurry signal were deleted, and this new version of the mesh was saved separately. The fibril directions were saved as a cell axis and added to the attribute map. Then the principal directions of growth were loaded onto the new mesh and the angle between growth directions and fibril directions was calculated. Then the distance heat map was loaded onto the new mesh, a proximal distal axis was created like before, and the angle between the fibril directions and the custom axis was calculated (which is the angle of the microtubules relative to the proximal-distal axis). Then to better visualize the microtubule signal at the surface of the image, the channel with the microtubule signal was loaded, and then the mesh was used to “annihilate” the signal everywhere except 1µm above and 1µm below the mesh. This annihilated image was used for screenshots of microtubule signal.

For the CATALASE inhibitor treatment (treatment details are described below), images were taken with a Leica Stellaris 5 using a 25X water dipping objective with an NA of 0.95 (HC FLUOTAR L VISIR 25X/0.95 WATER). A 488 laser with a power of 1 and a gain of 75, detecting wavelength of 494-550, was used to image GFP-MBD. A 561 laser with a power of 0.2 and a gain of 100, detecting wavelength of 582-607, was used to image propidium iodide. Channels were imaged in the same track, with line averaging of 2 and a scanning speed of 400, and bidirectional scanning. Images were a format of 1024x1024 pixels and 8 bit. Voxel size was x=0.0865799µm, y=0.0865799µm, z=0.2µm.

MorphoGraphX was used to process the images. A neural network was run on the propidium iodide channel to predict locations of the cell wall, which creates a stack with a brighter and more continuous surface. This new stack was then used with an edge detect signal threshold to detect the surface. This was used to create the mesh, and then the signal from the propidium iodide was projected onto the mesh and used for segmentation. Then to better visualize the microtubule signal at the surface of the image, the channel with the microtubule signal was loaded, and then the mesh was used to “annihilate” the signal everywhere except 1µm above and 1µm below the mesh and used for screenshots of microtubule signal. For the propyzamide treatments, plants with the *pUBQ10::GFP-MBD* transgene were dissected two days before imaging and placed on media. Images were taken with a Zeiss LSM 710 using a 20× water dipping objective with an NA of 1.0 (W Plan-APOCHROMAT20×.1.0DIC(US)VIS-IR). A 488 laser with a power of 25 and a gain of 750, detecting wavelength of 493-586, was used to image GFP-MBD. A 594 laser with a power of 2 and a gain of 670, detecting wavelengths of 599-641, was used to image propidium iodide. Channels were imaged in the same track, with line averaging of 4. Images were a format of 512x512 pixels and 16 bit. Voxel size was x=0.1383776, y=0.1383776, z=0.2 µm. For image processing, signal from propidium iodide is used with a neural network to predict the location of cell walls, and then the prediction is used with an edge detect signal threshold to detect the surface. This is used to create the mesh, and then the signal from the neural network predict is projected onto the mesh and used for segmentation. To better visualize the microtubule signal at the surface of the image, the channel with the microtubule signal was loaded, and then the mesh was used to “annihilate” the signal everywhere except 1µm above and 1µm below the mesh. Then the before and after treatment time points were lineage tracked, and cells are scored based on the microtubule signal.

For the 5 min time lapse imaging of microtubules, plants with the transgenes *pUBQ10::GFP-MBD* and *pUBQ10::mCherry-RCI2A* were dissected two days before imaging. Images were taken with a Leica Stellaris 5 using a 25X water dipping objective with an NA of 0.95 (HC FLUOTAR L VISIR 25X/0.95 WATER). A resonant scanner with a speed of 8000 was used with bidirectional scanning and line averaging of 16. A 488 laser with a power of 3 and a gain of 150, detecting wavelengths of 494-550, was used to image GFP-MBD. Images were a format of 1024x1024 pixels and 8 bit. Voxel size was x=0.0270563µm, y=0.0270563µm, z=0.2 µm for the stage 5 (earlier stage) sepals, and x=0.432899µm, y=0.432899µm, z=0.2 µm for the stage 7 (later stage) sepals. Images were set up and taken every 5 min for 65 min, and junctions between cells were used as landmarks to align images as best as possible. In ImageJ, max projections were combined into a stack, and then the plugin HyperStack (rigid body) was used to register the images. After registration, the time points were split into images, and the 0 min and 5 min time points were merged.

### Analysis of temporal and spatial heterogeneity (patchiness)

Analysis of heterogeneity was done as described in Burda et al 2024.

### Quantification of microtubule organization on a subcellular scale

Ilastik was used to classify microtubules as organized or crisscrossed. The images used were screenshots of ‘annihilated’ microtubule signal (described above), and the background and any blurred signal at the periphery was cropped out of the images. For Figure 3, a small subset of pixels on a few images in the data set were used for training, and then the rest of the pixels were batch processed. Although not completely unbiased, it is a way to classify visual impressions blindly and reproducibly. For Figure 4, the images were batch processed using the same classification system from Figure 3.

### CATALASE inhibitor treatment

Inflorescences with the microtubule marker *pUBQ10::GFP-MBD* were dipped in 200 µm 3-AT in a HEPES buffer with 0.02% Silwet and pH of 7 or a mock treatment of just HEPES buffer with 0.02% Silwet and pH of 7 once per day for 4 days. On day 5, inflorescences were either imaged or stained for hydrogen peroxide. For the inflorescences that were imaged, they were dissected, placed on apex culture media (described above) in a tilted position with flowers of about stage 6 at the top stained with 1.87 mM propidium iodide for 4 min by filling the petri dish with the solution with the inflorescences mounted in media, then the stain was removed, and inflorescences were rinsed twice by placing water on top of the inflorescences in the media. Images were taken 5-10 min after the last rinse, which is enough time for the position of the sample to stop moving. This process was performed one at a time for each inflorescence to minimize effects of ROS created by dissection. For the inflorescences stained for hydrogen peroxide, the DAB stain solution was made as described above, and inflorescences were stained for 25 hrs.

### Propyzamide treatment

For the propyzamide treatment, propyzamide was added to the apex culture media (described above) at the same time as the vitamins are added to the media. A concentrated stock of 10mM of propyzamide in DMSO was made, and then added to the media to create a 200µM final concentration. Propyzamide concentrations of 20µM, 100µM, and 200µM were tested. Media with 200µM propyzamide often caused complete depolymerization of microtubules after 24 hours. Sepals on 200µM propyzamide media were imaged once every 30 min, starting at 15 min after being moved to the media. A noticeable amount of depolymerization occurred between 15 min and 45 min. Between 45 min and 75 min the microtubules became short and thick, which could indicate some rate of polymerization. From 75 min to 225 min the rate of depolymerization was slower, which may again indicate some rate of polymerization. Therefore, a 30min treatment of 200µM propyzamide was chosen as optimal for causing depolymerization with minimal effects of re-polymerization.

Plants were dissected two days before the experiment to allow for recovery from dissection. One at a time, inflorescences were stained in 1.87 mM propidium iodide with 0.02% Silwet for 4 min in a lid of an Eppendorf tube, then rinsed by being placed in the lid of an Eppendorf tube with water, placed in normal apex culture media, and placed in a tilted position so that a stage 5-6 sepal was most visible. Then water was added to the petri dish for about 10 min before imaging to allow the inflorescence to settle into the media and water. Sepals were imaged, moved to propyzamide media in the same tilted position, water was added to the petri dish again, and imaged for a second time after 30 min on the propyzamide media.

### Oryzalin treatment

For the oryzalin treatment, oryzalin was added to the apex culture media (described above) at the same time as the vitamins are added to the media. A concentrated stock of 84.2mM of oryzalin in DMSO was made, and then added to the media to create a 100 µM final concentration. This concentration of oryzalin was enough to begin depolymerizing microtubules within 24 hours, and completely depolymerize microtubules within 48 hours. Higher concentrations precipitated in the media. Plants were dissected and then placed on normal media for one day to recover from dissection. After recovering from dissection for one day, inflorescences were moved to media with oryzalin or a DMSO mock treatment (and imaged on the third day, about 48 hours after dissection). Plants were moved to new media halfway through the imaging series.

### Statistical analysis

Pairwise p values for the temporal heterogeneity and patchiness scores in Figure 1 and 7 were obtained using z-test for correlation coefficients. Figure 3 p values were obtained using a T-test between genotype pairs using data from all replicates and days.

### Resource Availability

Lead contact: Requests for further information and resources should be directed to and will be fulfilled by the lead contact, Adrienne H.K. Roeder.

Materials Availability: The plasmid for *CAT2oe* and seeds for plant lines are available upon request to the lead contact without restriction.

## Data and Code Availability

● All data from live imaging series has been deposited at 10.5281/zenodo.15318675
● Code will be deposited at later submission
● Any additional information required to reanalyze the data reported in this paper is available from the lead contact upon request.

## Acknowledgments

We thank Byron Rusnak, Michelle Heeney, Lilijana Oliver, and Si Chen for helpful comments on the manuscript. We thank Arezki Boudaoud for conversations and feedback. We thank Ram Dixit for discussing concepts and protocols for depolymerizing microtubules. Research reported in this publication was supported by the National Institute of General Medical Sciences of the National Institutes of Health under award number R01GM134037 (to A.H.K.R.) and Gordon and Betty Moore Foundation post-doctoral fellowship award #2919 (to F.B.). The content is solely the responsibility of the authors and does not represent the views of the National Institutes of Health and other funders.

## Author Contributions

Conceptualization: IB, LH, AHKR. Data curation: IB. Formal analysis: IB, FB. Funding acquisition: AHKR. Investigation: IB, ES, LH. Methodology: IB, AS. Project administration: AHKR. Supervision: AHKR. Visualization: IB. Writing-original draft: IB. Writing-review and editing: IB, FB, AS, ES, LH, AHKR.

## Supplemental information titles and legends

**Figure S1:** ROS inhibits growth heterogeneity, related to Figure 1

**Figure S2:** Microtubule arrangement is different in *ftsh4-5,* and rescued in *ftsh4-5 CAT2oe*, related to Figure 2

**Figure S3:** ROS is sufficient to lead to crisscrossed microtubules, related to Figure 4

**Figure S4:** Microtubules in *ftsh4-5* are more stable, related to Figure 5

**Figure S5:** Depolymerizing the microtubules is insufficient to restore growth heterogeneity, related to Figure 7

**Figure S6:** Oryzalin treatment is effective in depolymerizing microtubules, related to Figure 7

**Video 1**, related to Figure 5

**Video 2**, related to Figure 5

