## Supplemental Figures for "ROS inhibits microtubule dynamics and cell growth heterogeneity during Arabidopsis sepal morphogenesis"

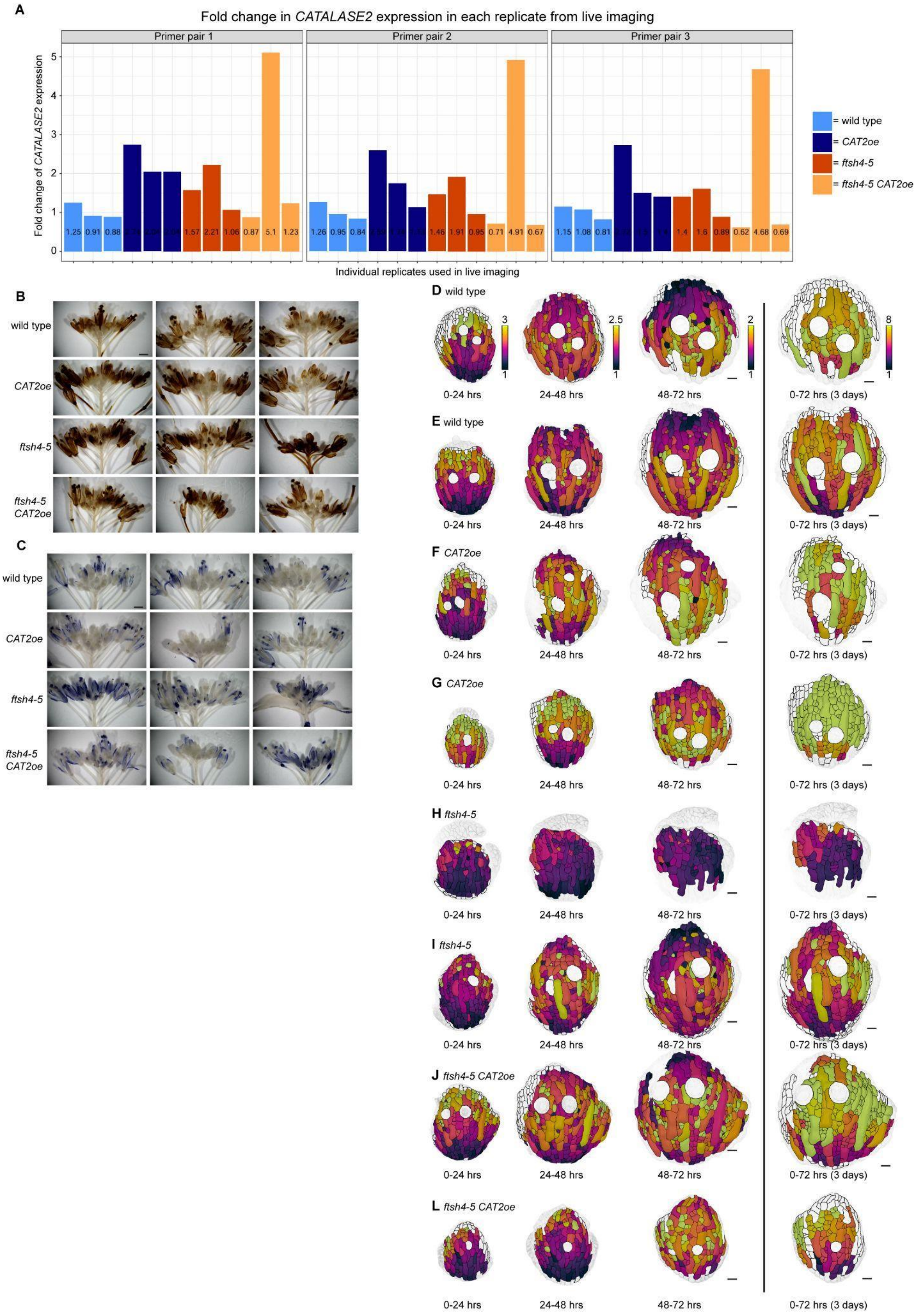
A
Fold change in CATALASE2 expression in each replicate from live imaging
Primer pair 1
Primer pair 2
Primer pair 3
Fold change of CATALASE2 expression
Individual replicates used in live imaging
= wild type
= CAT2oe
= ftsh4-5
= ftsh4-5 CAT2oe
B
C
wild type
CAT2oe
ftsh4-5
ftsh4-5 CAT2oe
D wild type
E wild type
F CAT2oe
G CAT2oe
H ftsh4-5
I ftsh4-5
J ftsh4-5 CAT2oe
L ftsh4-5 CAT2oe
0-24 hrs
24-48 hrs
48-72 hrs
0-72 hrs (3 days)

Figure S1: **ROS inhibits growth heterogeneity.** (A) Fold change in *CATALASE2* expression from QPCR of tissue from the same plants used for individual live imaging replicates. (B) Inflorescences stained for hydrogen peroxide. Scale bars are 0.5 mm. (C) Inflorescences stained for superoxide. Scale bars are 0.5 mm. I-L: Remaining replicates of live imaging. Cell area growth represented as a ratio, projected on the later time points over 24-hour intervals and cumulative over 3 days for (D-E) wild type, (H-I) *CAT2oe*, (J-K) *ftsh4-5*, and (L-M) *ftsh4-5 CAT2*. Scale bars are 20µm. Related to Figure 1.

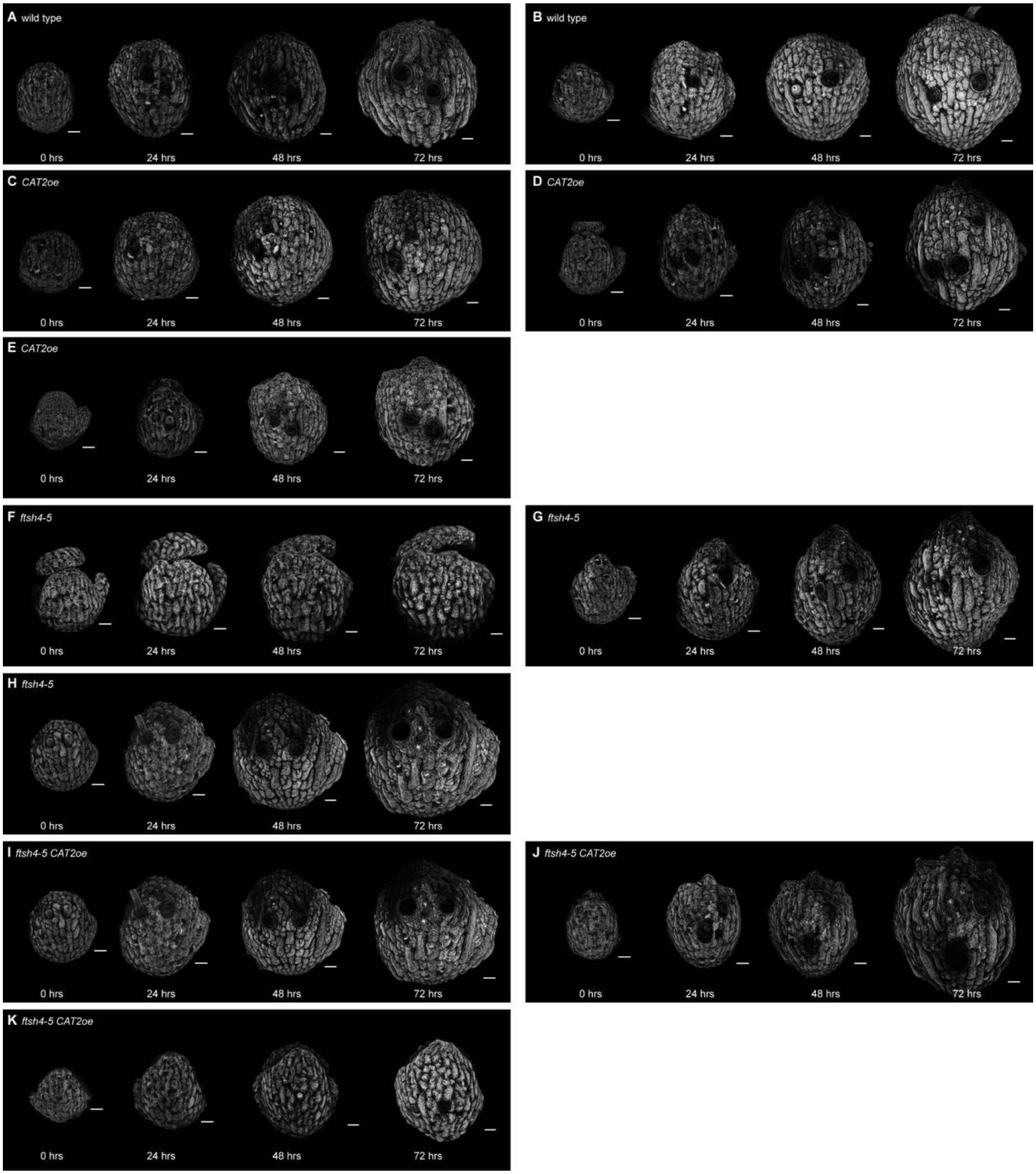


Figure S2: **Microtubule arrangement is different in *ftsh4-5* and rescued in *ftsh4-5 CAT2oe*.** A-J: Microtubule signal from the live time lapse imaging of sepal development. Other replicates of wild type (A-B), all three replicates of *CAT2oe* (C-E), other replicates of *ftsh4-5* (F-G), and all three replicates of *ftsh4-5 CAT2oe* (H-J). Scale bar is 20μm. Related to Figure 2.

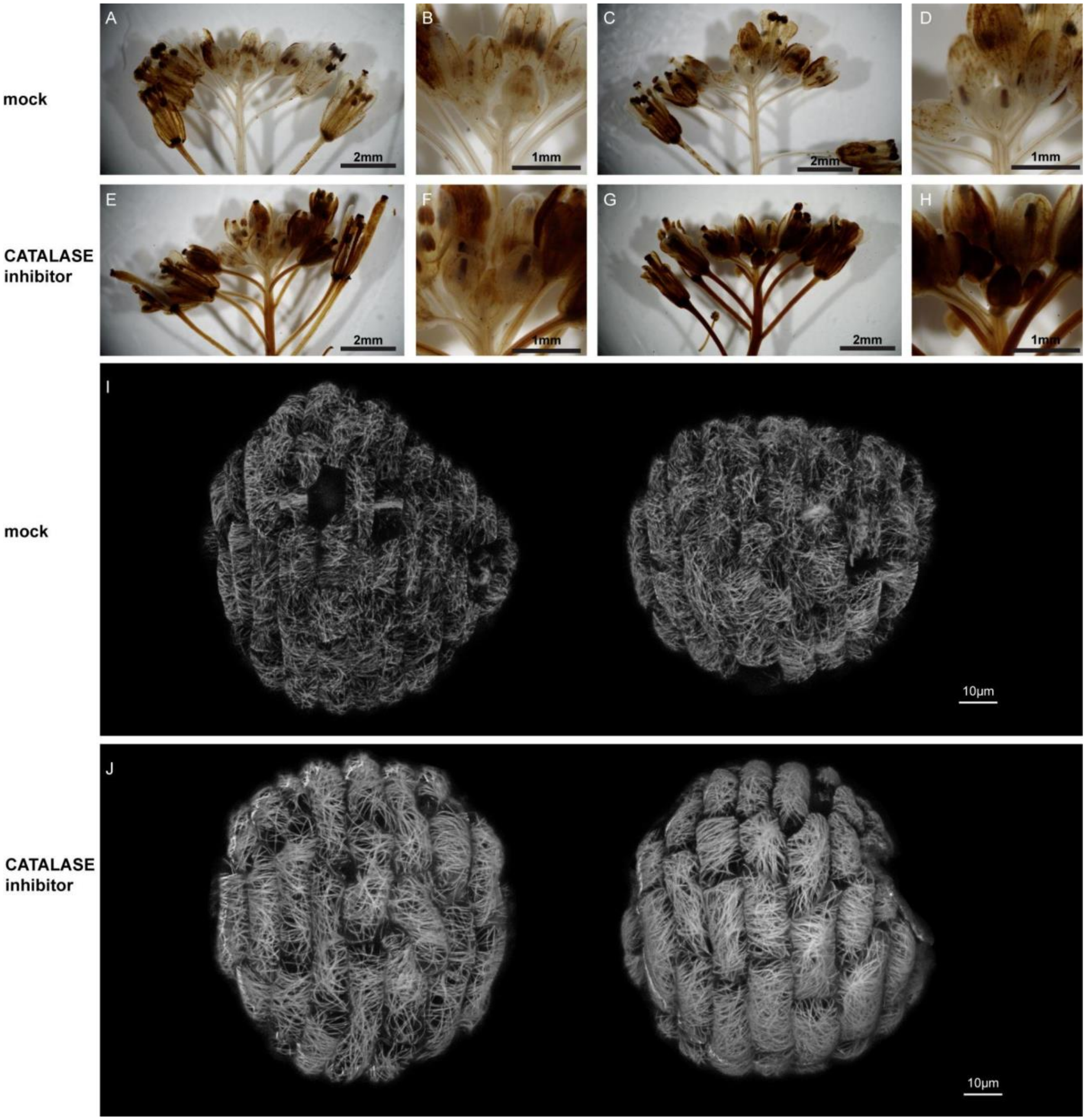


Figure S3: **ROS is sufficient to lead to crisscrossed microtubules.** A-:H Inflorescences are stained for hydrogen peroxide with DAB after a mock treatment (A-D) or a CATALASE inhibitor treatment (E-H). Other replicates are shown here. I-J: Microtubules in (I) mock-treated and (J) CATALASE-inhibitor treated sepals. Other replicates are shown here. Related to Figure 4.

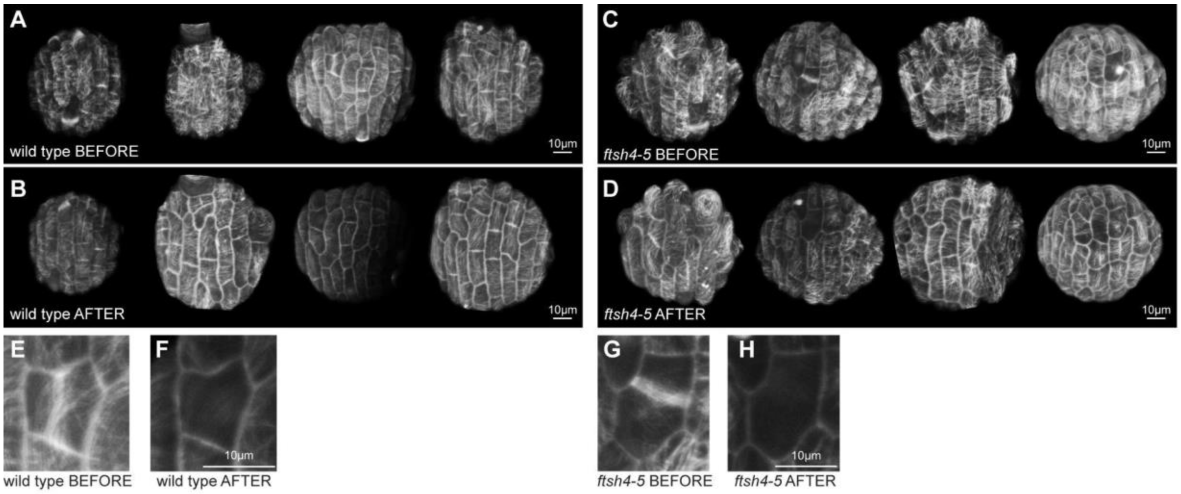


Figure S4: **Microtubules in *ftsh4-5* are more stable.** A-H: Other replicates of the propyzamide treatment to compare microtubule stability. (A) Wild type sepal cells before treatment and (B) the same wild type sepals  after 30 min propyzamide treatment. (C) *ftsh4-5* sepal cells before treatment, and (D) the same *ftsh4-5* sepals after 30 min propyzamide treatment. E-H: Zoomed in to show that microtubules forming the preprophase band depolymerize in both wild type (E-F) and *ftsh4-5* (G-H) indicating that the treatment is effective in both genotypes. Related to Figure 5.

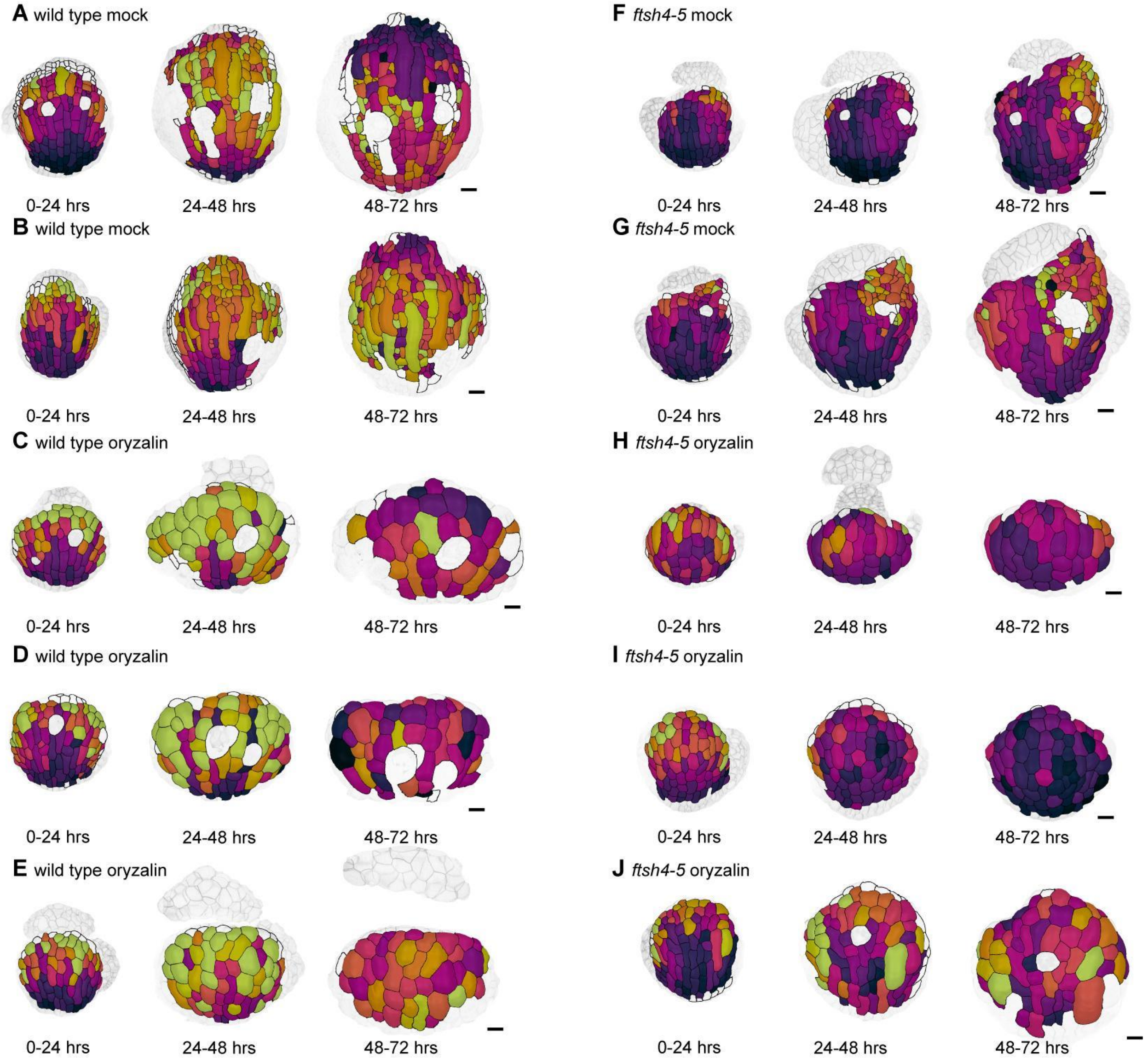


Figure S5:**Depolymerizing the microtubules is insufficient to restore growth heterogeneity.** Cell area growth heat maps of sepal development with an oryzalin treatment. Sepals were imaged once every 24 hours for 4 days. Area growth is represented as a ratio and projected onto the later time point. The rest of the replicates are shown for (A-B) wild type mock treatment, (C-E) wild type oryzalin treatment, (F-G) *ftsh4-5* mock treatment, (H-J) *ftsh4-5* oryzalin treatment. Related to Figure 6.

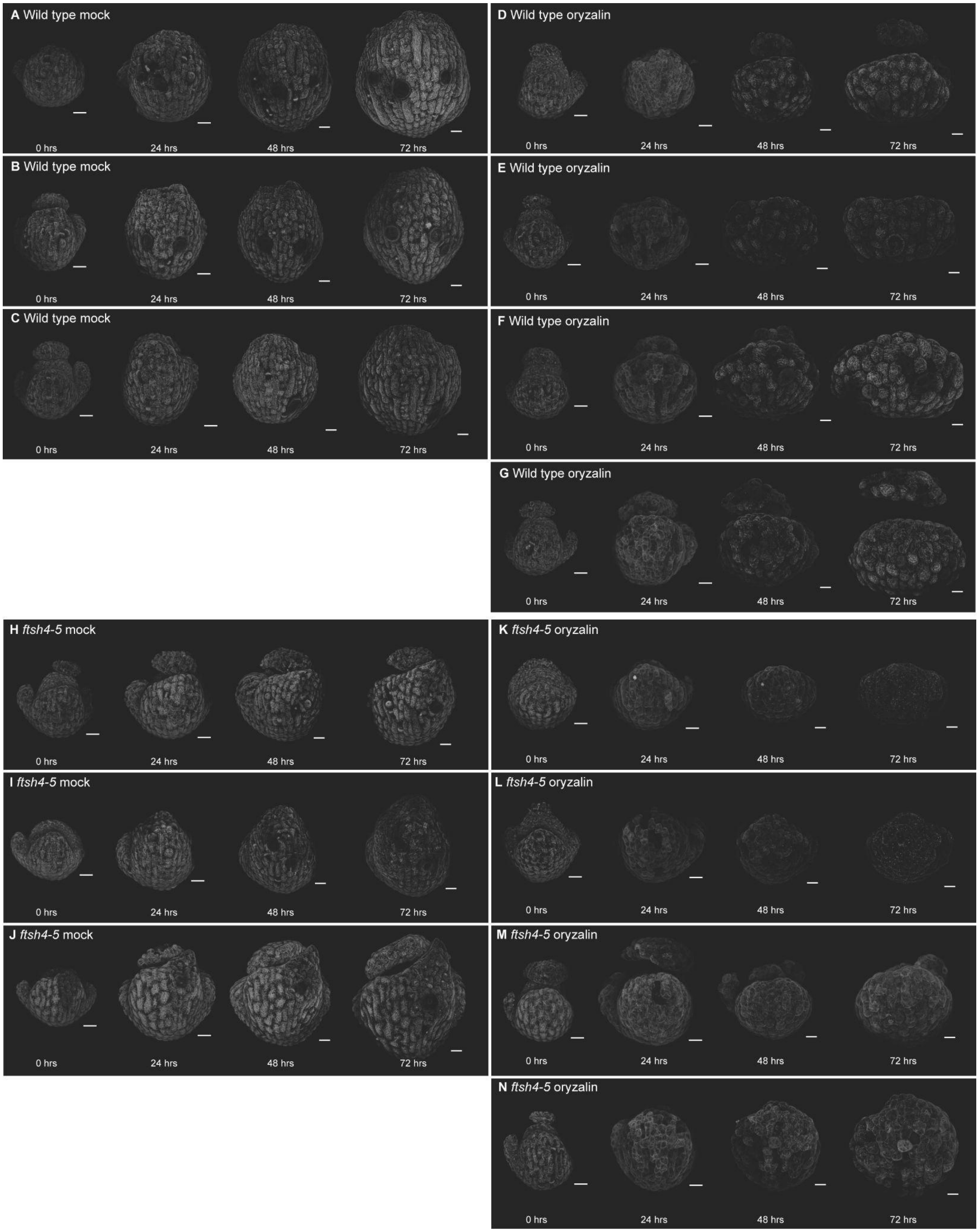


Figure S6: **Oryzalin treatment is effective in depolymerizing microtubules.** Microtubule signal from the live imaging with the oryzalin treatment. All replicates are shown for wild type mock (A-C), wild type oryzalin (D-G), *ftsh4-5* mock (H-J), *ftsh4-5* oryzalin (K-N). Related to Figure 6.
